# CBKMR: A Copula-Based Bayesian Kernel Machine Regression Framework for Optimal Marker Detection in Omics Data

**DOI:** 10.1101/2025.11.18.689140

**Authors:** Anirban Chakraborty, Chloe Mattila, Debashis Ghosh, Brian Neelon, Souvik Seal

**Affiliations:** Department of Public Health Sciences, Medical University of South Carolina; Department of Biostatistics and Informatics, Colorado School of Public Health

**Keywords:** Bayesian variable selection, GKMR, Gaussian Copula, nearest neighbor GP approximation, Bulk omics, scRNA-seq

## Abstract

High-throughput bulk and single-cell omics technologies enable comprehensive molecular profiling; yet identifying compact, biologically interpretable marker sets that distinguish cell types, conditions, or disease states remains challenging. Standard pipelines rely on univariate differential expression tests, which ignore gene–gene dependencies and nonlinear effects, while multivariate machine-learning (ML) methods often lack principled feature selection and uncertainty quantification. The Bayesian kernel machine regression (BKMR) framework offers an appealing alternative because it (a) captures nonlinear gene–outcome relationships and higher-order interactions, and (b) enables automatic relevance determination (ARD) through sparsity-inducing priors. We show that the traditional latent Gaussian process (GP) formulation of BKMR is inadequate for discrete outcomes (e.g., cell-type labels), leading to biased inference and unstable variable selection. Instead, we propose a copula-based Bayesian kernel machine regression (CBKMR) model that uses outcome-appropriate discrete marginals while a Gaussian copula captures kernel-induced dependence across observations. To ensure scalability to modern single-cell datasets, we further introduce a nearest-neighbor GP-based variant, NNCBKMR, which reduces computational complexity from *O*(*N*^3^) to nearly linear in *N*. Simulation studies show that CBKMR more accurately captures nonlinear effects and yields stronger marker-selection performance than BKMR and top ensemble ML methods (e.g., random forests, XGBoost). Applications to multiple scRNA-seq datasets demonstrate that CBKMR identifies concise marker panels that align closely with expert-annotated gene signatures while providingposterior uncertainty for principled decision-making.

## 1. Introduction

Over the past several years, high-throughput omics technologies, spanning both bulk and single-cell assays, have revolutionized our ability to quantify diverse molecular activities across complex biological systems (Wang and Bodovitz (2010); Dai and Shen (2022)). These platforms enable comprehensive profiling of genes, proteins, and metabolites at multiple levels of biological resolution, opening new avenues for exploring cellular heterogeneity, tissue organization, developmental trajectories, and disease mechanisms. A parsimonious yet robust panel of biologically meaningful markers, i.e., molecular features that consistently characterize specific cell types, conditions, or disease states, is essential for clinical translation, as these can stratify patients, guide diagnostic and prognostic decisions, and inform therapeutic targeting (Ghosh and Poisson (2009); Abu-Asab et al. (2011); Quezada et al. (2017); Olivier et al. (2019)). Such a panel is also crucial for de-signing next-generation spatial transcriptomics (ST) and proteomics studies, where costs are substantial and informed gene or protein selection can markedly reduce assay burden (Yafi et al. (2024)). Molecules often interact and exhibit nonlinear associations with biological outcomes (e.g., cell type identity or disease status), yet most analytical pipelines still rely on univariate differential expression (DE) tests (e.g., Anders and Huber (2010); Love, Huber and Anders (2014a); Costa-Silva, Domingues and Lopes (2017)) for ad hoc marker selection, which overlook these complexities and fail to produce concise, predictive, and clinically translatable marker panels (Pullin and McCarthy (2024)).

Standard software packages for single-cell RNA sequencing (scRNA-seq) analysis, such as Seurat (Hao et al. (2024)) and Scanpy (Wolf, Angerer and Theis (2018)), offer multiple options for detecting differentially expressed genes. These range from simple parametric and non-parametric pairwise tests, such as Welch’s t-test and the Wilcoxon rank-sum test, to more advanced model-based approaches, including hurdle model-based MAST (Finak et al. (2015)) and generalized linear models (GLMs) with negative binomial or Poisson likelihoods (Hilbe (2011)). However, such independent, per-gene testing frameworks fail to account for joint dependencies among genes, making it difficult to systematically define a minimal subset of genes that captures maximal biological variation. In practice, detected genes are typically ranked by false discovery rate (FDR)–adjusted *p*-values, and an arbitrary cutoff is then applied to select the top-ranking genes as marker genes, a procedure that is inherently subjective and seldom reflects the true multivariate structure of the data. Furthermore, these approaches may overlook nonlinear relationships between gene expression and biological outcomes. To address these limitations, recent efforts have increasingly focused on predictive, multivariate modeling using machine learning (ML) and deep learning (DL) frameworks (e.g., Aevermann et al. (2021); Dai, Pei and Wang (2022); Nelson, Riva and Cvejic (2022); Covert et al. (2023); Tangherloni et al. (2023); Gregory et al. (2024)). Some approaches leverage tree-based ensemble methods, such as random forests (Breiman (2001)) and XGBoost (Chen and Guestrin (2016)), to model cell type or condition (e.g., one-vs-rest) as a binary response and gene-expression profiles as predictors. However, determining predictor significance through feature-importance thresholding remains ad hoc and non-trivial. Other approaches adopt optimization-based algorithms, including DL architectures, which often require extensive hyperparameter tuning and careful regularization. Therefore, there is a need for intuitive, automated, and uncertainty-aware marker-detection methods that can be readily adopted in biological and clinical research.

With the rapid surge of genome-wide association studies (GWAS) two decades ago, kernel machine regression (KMR) frameworks (Cristianini and Shawe-Taylor (2000); Seeger (2004)) gained prominence as powerful tools for detecting nonlinear associations between individual or groups of SNPs and complex phenotypes (e.g., Liu, Lin and Ghosh (2007); Wu et al. (2011); Maity, Sullivan and Tzeng (2012); Larson and Schaid (2013); Ge et al. (2015)) . Fundamentally, KMR extends traditional linear regression by relaxing the linearity assumption and introducing a semiparametric function, typically modeled as a realization from a Gaussian process (GP) with an appropriate kernel covariance function (Rasmussen and Williams (2006)). The kernel covariance structure captures the similarity between samples based on their feature expression profiles. Similar models have been recently used in scRNA-seq datasets, facilitating analogous association testing between genes and phenotypic outcomes such as cell type identity or disease status (e.g., Zhang et al. (2022); Ozier-Lafontaine et al. (2024); Ghosh et al. (2025)). However, most of these studies emphasize the detection of nonlinearly differentially expressed genes, rather than predictive modeling aimed at identifying a compact, non-redundant set of informative markers. Consequently, the kernel covariance matrix is constructed using only a single feature or a small subset of features, implicitly assuming fixed or equal importance among them. In a conceptually related domain, namely, the identification of key exposures from complex environmental mixtures, the Bayesian KMR (BKMR) framework (Chakraborty et al. (2007); Bobb et al. (2015)) introduced a more flexible approach by leveraging the automatic relevance determination (ARD) kernel (Tipping (2001)), which allows predictors to assume varying levels of importance. By further placing a sparsity-inducing spike-and-slab prior (Ishwaran and Rao (2005)) on the inverse lengthscales, which govern the relevance of individual features, this formulation adaptively learns the relative importance of each predictor dimension within a unified modeling framework. The appeal of such an approach in our context is evident. One can envision a Bayesian logistic KMR (Mallick, Ghosh and Ghosh (2005)) with cell type or disease status as the outcome, where all genes are embedded within an ARD kernel covariance; the model automatically infers each gene’s relative importance and summarizes it through the posterior inclusion probability (PIP), providing an intuitive mechanism for selecting the final subset of informative markers.

Bobb et al. (2018) implemented a probit KMR model for binary outcomes with a similar variable selection strategy. More broadly, the KMR framework naturally extends to accommodate a wide class of exponential family outcomes, such as Gaussian, binomial, Poisson, and negative binomial, through the generalized KMR formulation (Ge, Smoller and Sabuncu (2016)). Mou, Zhang and Arshad (2025) recently introduced a Bayesian extension, generalized Bayesian KMR (GBKMR), which incorporates variable selection analogous to BKMR. These generalized KMR models share a common assumption: kernel-induced dependence is introduced through an *additive* latent Gaussian random effect inside the link function, implicitly requiring that the joint dependence among responses be fully captured by correlations in a multivariate Gaussian latent space. While appropriate for continuous outcomes, this assumption becomes restrictive for discrete or otherwise non-Gaussian data. As Mad-sen (2009) showed, for discrete outcomes the latent Gaussian correlation structure no longer *uniquely* determines the dependence among the observed responses. Moreover, the posterior inference for such methods generally require Laplace approximation of the likelihood, which can produce suboptimal estimation of the model parameters (Joe, 2008). Consequently, latent GP-based mixed models often underrepresent or distort the true dependencies across observations, particularly when the discrete outcome distribution is skewed or overdispersed. The above limitations motivate a copula-based construction (Sklar (1959)), which models the outcome distribution directly while using a flexible copula to encode kernel-driven dependence, thereby more accurately capturing complex, nonlinear, and non-Gaussian dependencies among discrete observations. We term the resulting framework *copula-based Bayesian kernel machine regression (CBKMR)*. Copula-based mixed models have a strong precedent in spatial statistics (Hughes (2015); Krupskii and Genton (2019); Ma et al. (2017); Pearse, Gunawan and Cressie (2025)), including the foundational work of Madsen (2009). By modeling outcome distributions and dependence separately through a Gaussian copula, CBKMR provides a more faithful representation of of the true dependence structure among the out-comes and enables principled feature selection in high dimensions. In extensive evaluations, including settings with model misspecification, logistic CBKMR consistently outperforms GBKMR and popular univariate DE tests such as Wilcoxon rank-sum test, presto, DESeq2 in detecting true features and provides a more parsimonious set of markers in comparison to leading ML ensemble methods such as random forests and XGBoost. It further recovers compact, pathologist-validated marker sets in three independent scRNA-seq datasets. To ensure scalability to larger datasets, we incorporate a nearest-neighbor Gaussian process (NNGP) approximation (Datta et al. (2016); Finley et al. (2019)) within the copula formulation, yielding *NNCBKMR*, an efficient variant that retains CBKMR’s rigor while reducing computational complexity from *O*(*N* ^3^) to nearly linear in *N*, where *N* denotes the total number of cells.

The remainder of the paper is organized as follows. Section 2.1–2.2 reviews GBKMR in the context of general exponential family distributions. Section 2.3 introduces the proposed CBKMR framework and details posterior inference. Section 2.5 presents the NNGP approximation for large datasets. Section 3 reports results from simulation studies. Findings from two scRNA-seq applications ar reported in Sec 4. We conclude in Section 5. An efficient `GitHub` package implemented in `Rcpp` is available.

## 2. Methods

### 2.1. Brief review of popular BKMR models

Conceptually related to the support vector machine (SVM) (Cortes and Vapnik (1995)), kernel-based Gaussian process (GP) methods have a longer history (Kimeldorf and Wahba (1971)) and have been widely applied in statistics and machine learning, including the Bayesian formulations (e.g., Smola and Schölkopf (2003); Vermaak, Godsill and Doucet (2003); Mallick, Ghosh and Ghosh (2005); Van Gestel et al. (2006); Minnier et al. (2015)). For clarity, we refer to the framework introduced by Bobb et al. (2015) as BKMR throughout this manuscript. Since its introduction, BKMR has undergone substantial methodological extension and has been applied across diverse scientific domains. As a non-comprehensive overview of key developments, Crawford et al. (2018) introduced Bayesian approximate kernel regression, which approximates BKMR by replacing the full GP kernel with a random Fourier feature representation, yielding substantial computational gains. Liu et al. (2018) extended BKMR to accommodate both longitudinal and cross-sectional correlations. Teng et al. (2020) treated the kernel function as a random object, performing *automatic kernel selection* based on model evidence. Mutiso et al. (2024) adapted BKMR for negative binomial outcomes, illustrating its utility in modeling associations between social vulnerability and COVID-19 mortality in South Carolina. Smith et al. (2025) expanded BKMR to incorporate heteroskedastic error structures. Most recently, Mou, Zhang and Arshad (2025) introduced a generalized BKMR framework, which serves as the foundation for our proposal.

### 2.2. Generalized Bayesian kernel machine regression

Suppose there are *N* cells, and we focus on a binary *one-vs-rest* classification for a given cell type. For *i* = 1, …, *N*, let *Y*_*i*_ denote the random variable representing the cell type label of the *i*^th^ cell, and let *y*_*i*_ be its observed value. Thus, *Y*_*i*_ is the underlying stochastic outcome, while *y*_*i*_ is its realization in the data. For the *i*^th^ cell, we further consider two sets of covariates: **x**_*i*_ ∈ ℝ^*b*^, which is assumed to have a linear effect on *y*_*i*_ (e.g., batch indicators), and **z**_*i*_ ∈ ℝ^*p*^, which is assumed to have a potentially nonlinear effect on *y*_*i*_ (e.g., gene expression profiles). For a general discrete outcome, GBKMR assumes that *Y*_*i*_ follows an exponential family distribution with density 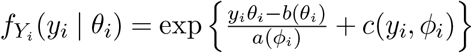, where *E*(*Y*_*i*_) = *µ*_*i*_, and the functions *a*(·), *b*(·), and *c*(·) are determined by the choice of the distribution for *Y*_*i*_. In the binary case *Y*_*i*_ ∈ {0, 1}, the model specializes to a Bernoulli likelihood with canonical logit link,

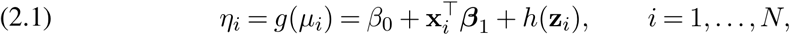

where *g*(·) = logit(·), *β*_0_ is the intercept, and ***β***_1_ is the coefficient vector for the linear covariates **x**_*i*_. The term *h* : ℝ^*p*^ → ℝ denotes a nonlinear function in a reproducing kernel Hilbert space (RKHS) H_*K*_ with a positive semidefinite reproducing kernel *K* : ℝ^*p*^ × ℝ^*p*^ → ℝ. Under this assumption, the vector of evaluations **h** = (*h*(**z**_1_), …, *h*(**z**_*N*_ ))^⊤^ follows an *N* -variate Gaussian distribution, **h** ∼ *N*_*N*_ (**0**, *τ* **K**) where *N*_*N*_ denotes an *N* -variate Gaussian distribution, *τ >* 0 is a scale (roughness) parameter, and 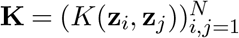 is the kernel covariance matrix constructed based on *K*. GBKMR adopts an automatic relevance determination (ARD) Gaussian kernel (Tipping (2001)), which assigns a separate *inverse* length-scale (relevance) parameter *r*_*m*_ ≥ 0 to each feature dimension:

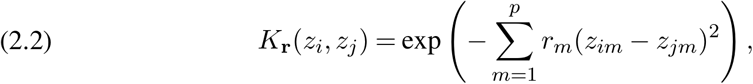

where **r** = (*r*_1_, …, *r*_*p*_)^⊤^. A large *r*_*m*_ indicates greater relevance of the *m*^th^ feature. The parameter set for (2.1)–(2.2) is 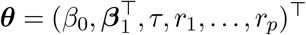, The Markov chain Monte Carlo (MCMC) procedure for posterior inference on ***θ*** based on an iterative reweighted least squares (IRLS) approach is provided in Mou, Zhang and Arshad (2025).

While embedding the additive random function *h*(·) inside *g*(·) in equation (2.1) offers a simple formulation for modeling the outcome, it is not guaranteed to recover the true dependence structure among observations. When *Y*_*i*_ follows a Gaussian marginal distribution and *g*(·) is the identity link, integrating *h*(·) yields the analytically exact joint distribution *N N* (*β*_0_**1** + **X*β***_1_, *τ* **Kr** + *σ*^2^**I**). However, as Madsen (2009) has noted, when the marginal distributions are non-Gaussian – as is the case for binary outcomes – such additive structure underestimates the true dependence among the *Y*_*i*_’s. Furthermore, integrating *h*(·) no longer yields a closed-form solution. Consequently, the inverse length-scales *r*_*m*_ cannot be straightforwardly interpreted as measures of overall gene-outcome association, limiting the biological interpretability of GBKMR for discrete outcomes.

### 2.3. Copula-based Bayesian kernel machine regression (CBKMR)

To construct the copula-based BKMR (CBKMR) model, we retain the previous notation and let *F*_*i*_(·) denote the cumulative distribution function (CDF) of *Y*_*i*_. As in (2.1), we adopt a logit link to specify the marginal mean structure, modeling *only* the linear effects,

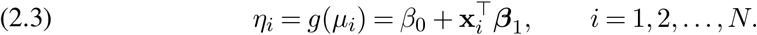

We focus on a Bernoulli outcome for clarity; however, the model is directly applicable to other discrete distributions through a suitable choice of link function *g*(·). We assume a Gaussian copula constructed from the marginal CDFs: 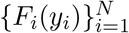 to characterize dependence among the observations *y*_*i*_ (Madsen (2009)), resulting

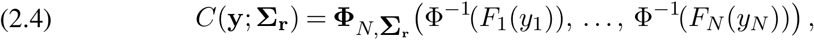

where *C* : ℝ^*N*^ → ℝ denotes the Gaussian copula CDF, 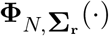 is the CDF of an *N* -variate normal distribution with correlation matrix **Σ**_**r**_, and Φ(·) is the CDF of a univariate standard normal distribution. We parameterize the correlation matrix as

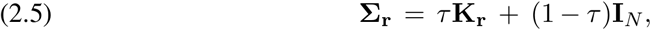

where 0 *< τ* ≤ 1 and **K**_**r**_ is the ARD Gaussian kernel from (2.2). The shrinkage parameter *τ* controls the contribution of nonlinear dependence: when *τ* ≈ 1, CBKMR behaves like a fully kernel-dependent model, whereas smaller *τ* values shrink **Σ**_**r**_ toward the identity, allowing observations to become (partially or fully) independent. This decomposition stabilizes inference in weak-signal settings and mitigates over-smoothing from the copula kernel.

Our next objective is to derive a density function corresponding to *C*(·). When F_i_’s are continuous, the corresponding density is uniquely defined on ℝ^*N*^ and can be easily derived using Sklar’s theorem (Nelsen (2006)),

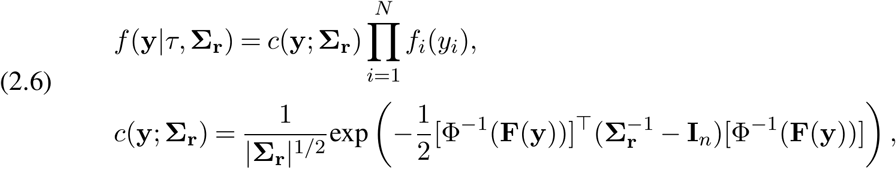

where **F**(**y**) = (*F*_1_(*y*_1_), …, *F*_*N*_ (*y*_*N*_ ))^⊤^ and *c*(**y**; **Σ**_**r**_) is the Gaussian copula density. However, because the marginals *F*_*i*_ of the discrete random variables *Y*_*i*_’s are step functions in our setting, the copula *C* does not possess continuous support on ℝ^*N*^, and Sklar’s theorem cannot be applied via differentiation to obtain a valid likelihood (Hughes (2015)). In principle, the likelihood could be recovered through an inclusion–exclusion expansion, but this requires summing over 2^*N*^ terms (Madsen (2009)), making it computationally infeasible even for moderate *N* . To overcome this limitation, we adopt a distributional transform (DT) approximation (Kazianka and Pilz (2010)), which has been shown to yield accurate Gaussian copula likelihoods for discrete marginals. For any *W*_*i*_ ∼ Unif(0, 1) independent of *Y*_*i*_, the DT of *Y*_*i*_ is defined as

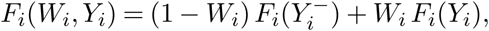

where 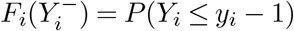. Following Hughes (2015), we replace *F*_*i*_(*y*_*i*_) by its conditional expectation under the DT,

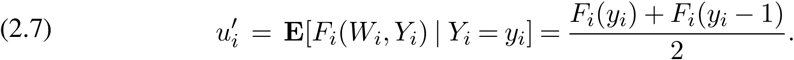

Substituting 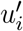 into the copula representation in (2.6) yields the approximate CBKMR like-lihood:

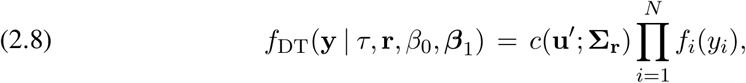

where 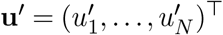, and 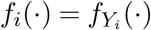 denotes the Bernoulli probability mass function (pmf). Such a copula-based BKMR formulation offers a key advantage over GBKMR: it is invariant to the choice of the marginal distribution. For any outcome type, we only specify a suitable link function *g*(·) to relate *µ*_*i*_ to **x**_*i*_ and define the marginals *F*_*i*_(·), while the dependence structure remains unchanged. Moreover, the *r*_*m*_s of CBKMR offers a clear biological interpretation for binary outcomes – if *r*_*m*_ *>* 0, it indicates that proximity along the *m*-th gene’s expression strongly correlates with cells sharing the same binary outcome. Thus, rather than interpreting a selected gene as merely shifting a baseline marginal probability, it is interpreted as defining the nonlinear decision boundary that separates the joint cell populations. Consequently, the *m*^th^ feature can be interpreted as a predictive biomarker – consistent with the definition of a predictive biomarker FDA-NIH BEST resource (Cagney et al., 2018). On the other hand, GBKMR embeds the kernel-induced dependence *h*(·) *inside* the link function. Not only this can be restrictive or unstable for discrete responses due to non-Gaussian dependence among observations (Joe, 2008; Madsen, 2009; Hughes, 2015), but such a construction also makes the interpretation of *r*_*m*_ difficult.

By separating marginal and dependence modeling, CBKMR provides more robust inference for discrete outcomes, as demonstrated in Section 3 through simulation studies. We next describe variable selection and the MCMC algorithm used to estimate the parameter vector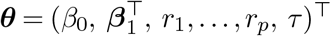

### 2.4. Sparsity prior and posterior inference

To devise an MCMC algorithm for ***θ*** we first assign a Beta prior distributions on the shrinkage parameter *τ* and Gaussian prior on the parameters *β*_0_ and ***β***_1_,

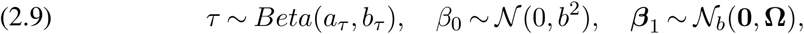

where *a*_*τ*_, *b*_*τ*_, *b*^2^ and **Ω** are prespecified hyperparameters. To induce sparsity within the ARD kernel and enable effective variable selection, the prior on the relevance parameters {*r*_*m*_} must place positive mass at zero, allowing weakly informative features to be *switched off*. Following Bobb et al. (2015), we consider a spike-and-slab prior (Ishwaran and Rao (2005)) as

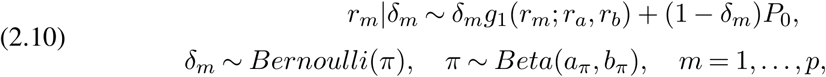

where *g*_1_(*r*_*m*_; *r*_*a*_, *r*_*b*_) is the density of a *Uniform*(*r*_*a*_, *r*_*b*_) distribution, *P*_0_ is a degenerate mass at 0, and *a*_*π*_ and *b*_*π*_ are the parameters of the Beta distribution. The full posterior distribution of the set of augmented parameters 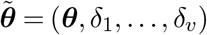 can be expressed as,

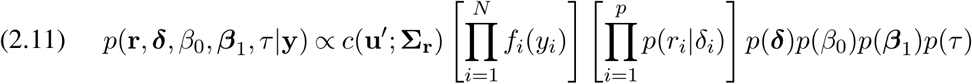

where *c*(**u**^′^, **Σ**_**r**_) is the copula density in equation (2.6), *p*(·)’s denote the prior densities of the corresponding parameters. Throughout the remaining of this paper, the value of the hyperparameter follows. First, we fix *a*_*τ*_ = *b*_*τ*_ = 1 that results in an uniform distribution for *τ* . The value of *b* is fixed at 1. For the slab counterpart in equation (2.10), we fix *r*_*a*_ = 0 and *r*_*b*_ = 3. Finally, to facilitate a parsimonious selection of markers in high dimensions, we fix *a*_*π*_ at 2.5 and set 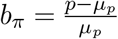, where *µ*_*p*_ = max(10, *p/*10). This creates a left skewed distribution for *π*, allowing the MCMC to converge faster. After fixing the hyperparameters, the MCMC algorithm proceeds in three steps - (i) update of *β*_0_ and ***β***_1_, (ii) update of the relevance and inclusion parameters (*r*_*m*_, *δ*_*m*_), and (iii) update of the shrinkage parameter *τ* . Unlike standard logistic GLMs, where Pólya–Gamma data augmentation (Pillow and Scott (2012); Polson, Scott and Windle (2013)) allows for direct sampling of *β*_0_ and ***β***_1_, the presence of the copula density *c*(·) in our model precludes standard Gibbs updates. Consequently, we implement a Metropolis-Hastings (MH) algorithm to sample these parameters, leveraging a modified Pólya–Gamma augmentation to facilitate efficient mixing. Update of *τ* is obtained through a random walk MH-based algorithm. Finally, we follow Bobb et al. (2015) to perform a reversible jump Markov chain Monte Carlo (RJMCMC) algorithm for updating **r** and ***δ***. We elaborate the steps in Algorithm 1. Once MCMC is completed and burn-ins are discarded, we calculate posterior inclusion probabilities (PIPs) for the *m*^th^ covariate in **z**_*i*_’s as 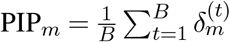. For a given threshold *thres* (say, 0.75), the *m*^th^ covariate is selected if PIP_*m*_ ≥ *thres*. A higher value of *thres* leads to a conservative selection of variables.

#### Algorithm 1

MCMC algorithm for CBKMR

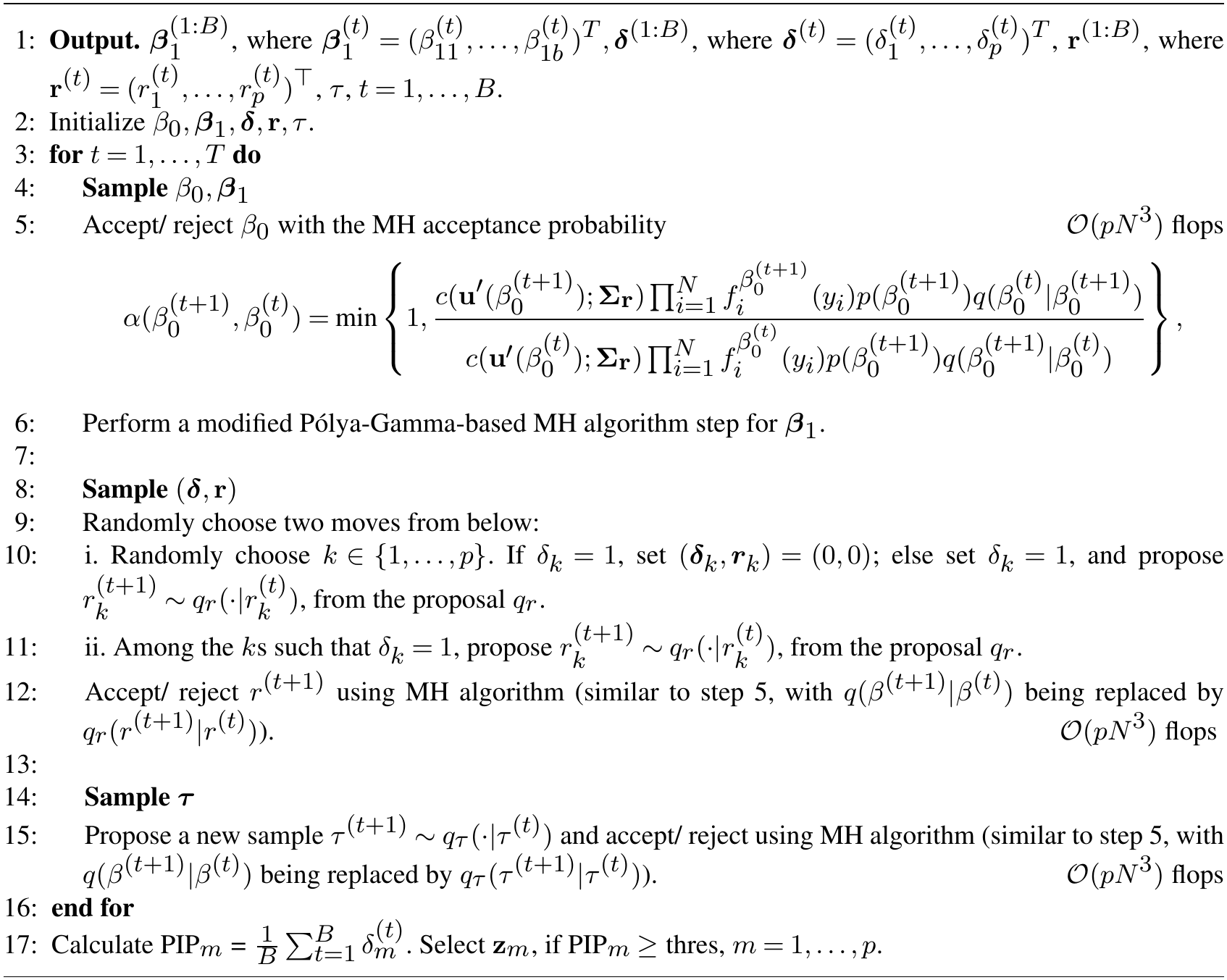

#### 2.4.1. CBKMR achieves superior variable selection performance compared to GBKMR

We now provide a mathematical justification for the superior variable selection performance of CBKMR over GBKMR observed in Section 3. To make the argument precise, we consider estimating a single inverse length-scale *r*_*m*_ by fixing the remaining *p* − 1 parameters at their true values 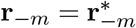 and 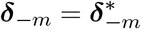, where 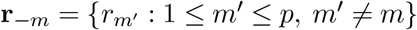 and ***δ***_−*m*_ follows the same convention. Under this reduction, the variable selection problem for feature *m* can be cast as the hypothesis test

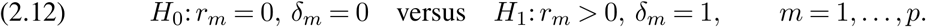

Both *H*_0_ and *H*_1_ carry the same interpretation under CBKMR and GBKMR: the null states that feature *m* contributes no signal to the kernel and is irrelevant, while the alternative asserts that it contributes a nonzero signal and should be selected. The binary selection decision *δ*_*m*_ is therefore *model-consistent*, providing common ground for comparing the two methods. We exploit a score-based test statistic to formalize this comparison.

##### Definition 2.1

(Score statistic and Fisher information). Consider the hypothesis testing framework in (2.12). For a method * ∈ {CBKMR, GBKMR}, let *f* ^*^(***θ***) denote the corresponding likelihood as a function of the parameter vector ***θ***. Define the score function for *r*_*m*_, evaluated at *r*_*m*_ = *r*, as

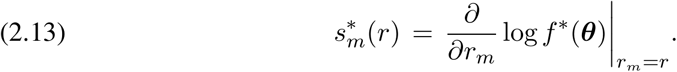

The Fisher information for *r*_*m*_ at *r*_*m*_ = *r*, under the null distribution of ***y***, is

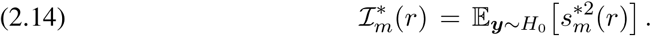

The standardised score statistic (Rao, 1948) is then defined as

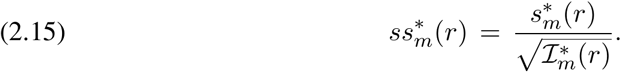

We now state a key lemma to establish our theoretical argument.

##### Lemma 2.2

(Fisher information ordering). *Under the null hypothesis in* (2.12), *the Fisher information satisfies*

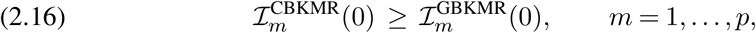

*where ℐ is as defined in Definition 2.1*.

Lemma 2.2 reveals that, at the null, the CBKMR likelihood is inherently more informative about the relevance parameter *r*_*m*_ than its GBKMR counterpart. Intuitively, this gain arises because the IRLS approximation underlying GBKMR inflates the effective working covariance beyond the true copula covariance of CBKMR, thereby lowering the precision with which *r*_*m*_ can be identified. Crucially, this information advantage is not confined to the null: since the noncentrality parameter – which governs power under the alternative – inherits its structure from the same Fisher information, the gain at *H*_0_ propagates directly into superior power at *H*_1_. The following theorem makes this connection precise.

##### Theorem 2.3

(Power superiority of CBKMR). *Consider the testing framework in* (2.12) *and suppose H*_1_ *holds, i*.*e*., *δ*_*m*_ = 1 *and* 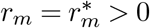. *The score test based on the CBKMR likelihood has a noncentrality parameter no smaller than that of the GBKMR IRLS-based score test:*

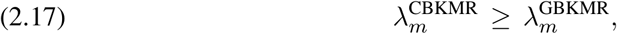

*where, for a method* 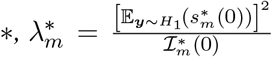, *with* 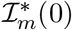 *as in Definition 2*.*1. Consequently, at every significance level α* ∈ (0, 1), *CBKMR achieves power no less than GBKMR:*

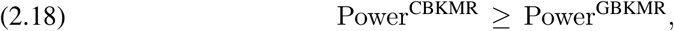

*where the power of the level-α score test is*

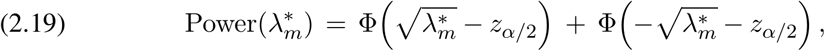

Φ(·) *is the standard normal CDF, z*_*α/*2_ = Φ^−1^(1 − *α/*2) *is the upper* (*α/*2)*-critical value, and the score statistic follows* 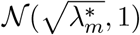 *under H*_1_ (*Demidenko, 2020*).

Since the power in (2.19) is strictly increasing in 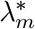, the inequality 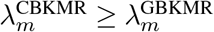 directly implies the power ordering at every *α* and *N* . To maintain the flow of the exposition, we defer the proofs of Lemma 2.2 and Theorem 2.3 to Appendix A.

#### 2.4.2. Prediction at new observation

After performing inference on existing observations, it is generally of interest to predict the unobserved response for a new sample with known covariates. Proposition 1 outlines the approximate conditional probability mass function of the unobserved response.

##### Proposition 1.

*Suppose* 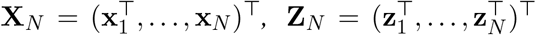 *and* **y** = (*y*_1_, …, *y*_*N*_ )^⊤^ *denote the observed covariates and responses for N individuals. The conditional predictive distribution of a random variable Y*_*N*+1_ *for a new observation with known covariates* **x**_*N*+1_ *and* **z**_*N*+1_, *given a specific set of parameter values, follows a discrete distribution with its probability mass function approximated as*

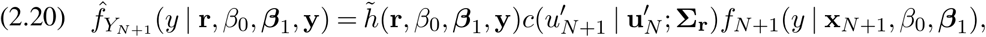

*where f*_*N*+1_(· | **x**_*N*+1_, *β*_0_, ***β***_1_) *denotes the marginal probability mass function of Y*_*N*+1_, *and* **u**_*N*_ *is the vector of latent normal transforms of the observed responses. The term c*(*u*_*N*+1_ | **u**_*N*_ ; **Σ**_**r**_) *is the conditional Gaussian copula density on the discrete transformed CDF’s of the old and new observations, i*.*e*., 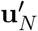 *and* 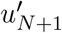 *respectively, and h*(·) *is a normalizing constant, such that*, 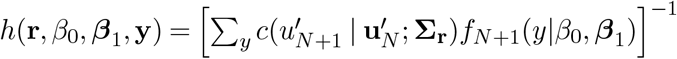.

When *Y*_*i*_ ∈ {0, 1}, we can easily compute the probability mass function using Proposition 1. Furthermore, by varying the values of the parameters, we can obtain a continuous distribution over the success probabilities.

### 2.5. Nearest neighbor CBKMR (NNCBKMR) for moderate p

The MCMC algorithm in Section 2.4 involves computation of the copula density *c*(·) in equation (2.8). This incurs *O*(*N* ^2^) memory and *O*(*N* ^3^) time at every iteration due to matrix-inversion of **Σ**_**r**_(Rasmussen and Williams (2006)). Consequently, CBKMR is prohibitive to implement even for a moderately large *N* (e.g., ∼ 2, 000) and *p* is of moderate size (say, 50). Here, we propose a scalable CBKMR approach by adopting a nearest neighbor Gaussian process (NNGP)-based sparse Cholesky decomposition approximation (Vecchia (1988); Datta et al. (2016); Katz-fuss and Guinness (2021)). To elucidate our approach, we temporarily suppress all the notations of CBKMR, and assume that a random vector **x** ∼ *N*_*N*_ (**0, Σ** = (**UU**^⊤^)^−1^), where **U** is the Cholesky factor of **Σ**^−1^. Motivated by the decomposition of any *N* -variate density into *N* univariate conditional densities 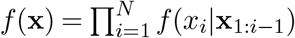, Vecchia (1988) proposed an approximate density 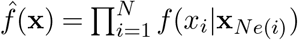, where *Ne*(*i*) ⊂ {1, …, *i*} is a subset of suitably chosen nearest neighbors (NNs) for the indices *i*. When applied to a *N* -variate Gaussian distribution, this yields a lower triangular sparse Cholesky factor **U**_S_ of **Σ**^−1^, such that **U**_S;*i,j*_ = 0 whenever *j* ∈*/ Ne*(*i*) (Datta et al. (2016); Katzfuss and Guinness (2021)). The sparsity structure *S* contains the information of neighbors *Ne*(*i*)’s. By fixing the maximum neighbor size max_*i*_|*Ne*(*i*)|= *k*, computation of the Gaussian likelihood using **U**_S_ requires only *O*(*Nk*) memory and *O*(*Nk*^3^) time. While *k* = *N* − 1 gives back the exact decomposition, in practice almost accurate approximation can be achieved with *k* ∼ 40 even for a very large *N* (Finley et al. (2019)). Since the copula density *c*(·) imitates a Gaussian likelihood, this sparse decomposition can be implemented to compute *c*(·).

#### Definition 2.4.

Suppose, **U**_**r**,*S*_ denotes a lower triangular sparse Cholesky factor of 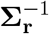 in (2.8) for a given sparsity structure *S*, such that max_*i*_|*Ne*(*i*)|= *k*. Then the NN approximated density of **y** is given by

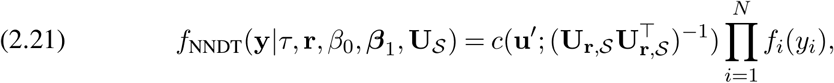

In addition to a guaranteed fast computation, *f*_NNDT_ is also approximately optimal in terms of Kullback-Leibler (KL) divergence minimization.

#### THEOREM 2.5.

*Suppose* **y** *is a N-variate vector that follows a Gaussian copula C*(**y**; **Σ**_**r**_) *with corresponding density c*(·), *continuous marginals* {*F*_*i*_} *and density* {*f*_*i*_}. *In addition, assume* **U**_**r**,S_ *is the sparse Cholesky decomposition of* 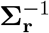 *for a given sparsity structure. Then*

i. *The nonzero entries in the i*^*th*^ *column of* **U**_**r**,*S*_ *are given by* 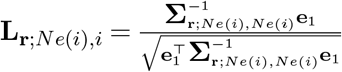,
ii. **U**_**r**,*S*_= argmin *KL*(*f*_*DT*_(**y**|***θ***)||*f*_*NNDT*_(**y**|***θ***, *A*_S_)), *A*_*S* _∈R_*N*×*N*_ :*A*_*S* _*>*0,*A*_*S,ij*_=0 ∀*j>i,j*∉^*N e*(*i*)^

*where* ***θ*** = {*τ*, **r**, *β*_0_, ***β***_1_}, **e**_1_ ∈ ℝ^|*Ne*(*i*)|×*i*^ *has the first entry equal to 1 and all other entries equal to 0. A*_S_ *>* 0 *implies that A*_S_ *is positive definite*.

To maintain clarity of the discussion, we postpone the proof to Section B in the appendix. The lemma 2.5 suggests that the NN approximated *f*_NNDT_(**y**|*τ*, **r**, *β*_0_, ***β***_1_, **L**_**r**,S_) is an optimal approximation to *f*_DT_(**y**|*τ*, **r**, *β*_0_, ***β***_1_), in Eq. 2.8.

To compute **U**_**r**,S_, we first perform a *maxmin* permutation (Guinness (2018)) of the observations and use a *k*-d tree-based algorithm from `GpGp` R package to find the nearest neighbors from the relevance-transformed inputs 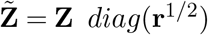. The resulting *f*_NNDT_ simply replaces *f*_DT_ in Section 2.4 for the MCMC steps of NNCBKMR. For clarity, we provide a pseudocode in algorithm 2.

#### Algorithm 2

MCMC algorithm for NNCBKMR

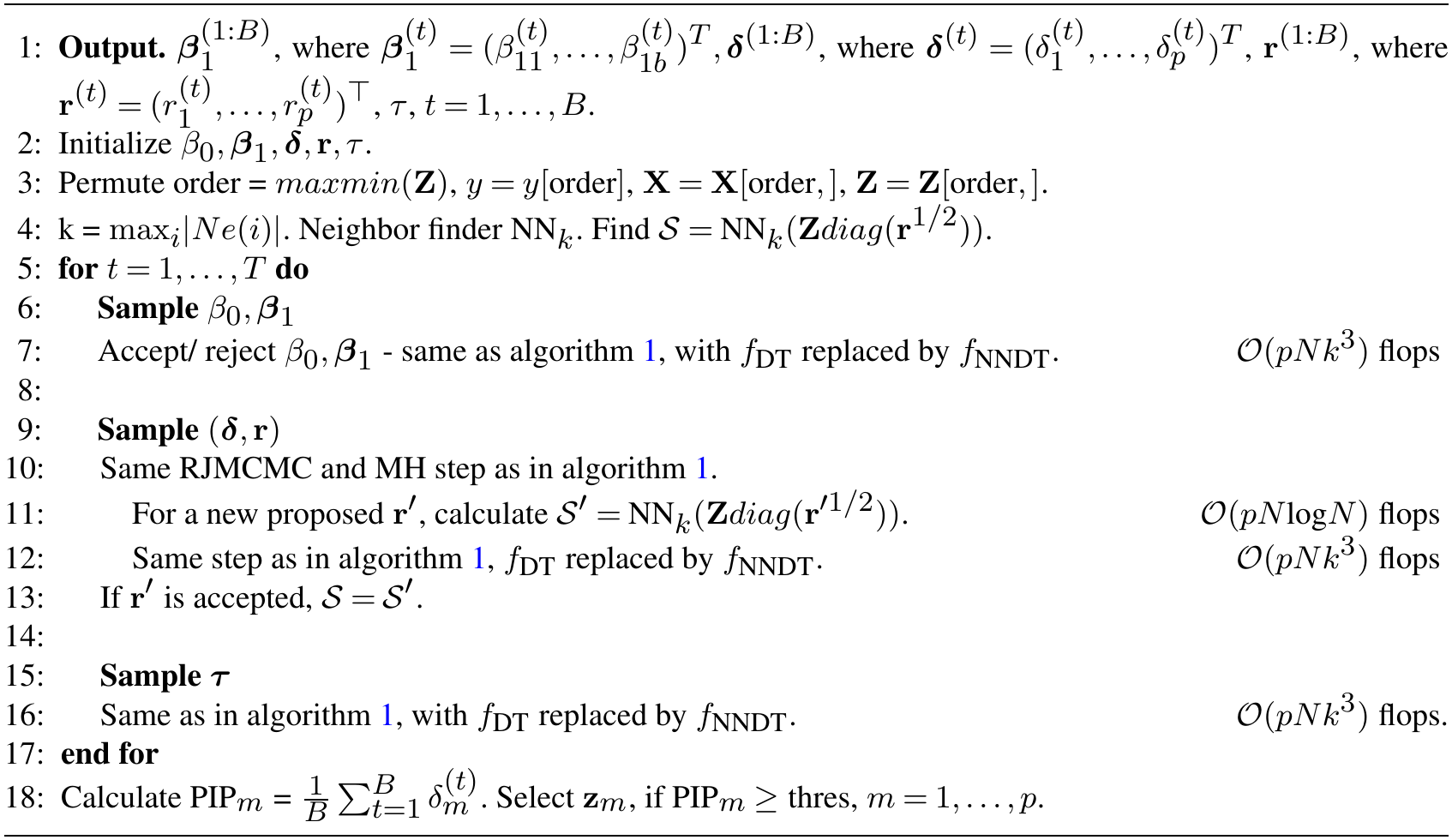

While NNCBKMR is computationally efficient when p is moderate, say in 50’s, its performance degrades significantly when *p* grows. This is due to the computational cost of finding the nearest neighbors that incurs a time complexity of *O*(*pN* log *N* ) (Friedman, Bentley and Finkel (1977)) that needs to be performed at every iteration following the update of **r**. Therefore, to apply NNCBKMR, we recommend reducing the dimension of the dataset using some FDR-based filtering.

## 3. Simulation experiments

Our simulation experiments share two common parameters *N* and *p*. To generate the design matrix **Z** = (**z**_·1_, …, **z**_·*p*_), where **z**_·*m*_ = (*z*_1*m*_, …, *z*_*Nm*_)^⊤^, we created two distinct simulation scenarios. In each of the simulation settings, We fixed *β*_0_ = 0 and ***β***_1_ = **0**, i.e., no intercept or linear covariates **x**. The detail of the scenarios follows.

### Scenario 1

To simulate **Z**, we first generated **z**_·1_ from an *N* -variate standard Gaussian distribution. For *m* = 2, …, *p*, we sampled an *N* -variate standard Gaussian vector and set **z**_·*m*_ to be the residual from regressing this vector on **z**_·1_, …, **z**_·*m*−1_. This yields mutually orthogonal covariate vectors while preserving Gaussianity. To assess robustness to different dependence structures, we considered two simulation designs in this scenario.

### Design 1.

We first simulated ***η*** = (*η*_1_, …, *η*_*N*_ ) from a *N* -variate Gaussian distribution,

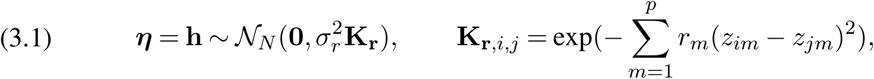

where 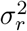 is the marginal variance and **r** = (*r*_1_, …, *r*_*p*_)^⊤^ is the p-variate relevance vector. We considered only the first 3 variables to be significant, i.e., *r*_*m*_ = 1, for *m* ≤ 3, and *r*_*m*_ = 0 ∀*m >* 3. By varying the values of *r*, 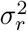, *N* and *p* we created several simulation scenarios under this setting. The marginal kernel variance 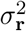 effectively controls how much of the relevances in **r** will be reflected in the response. A smaller value of 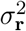 would eliminate any effect of **r**. As 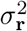 increases, the effect of **r** becomes more prominent.

### Design 2.

We considered another additive model but with more significant covariates,

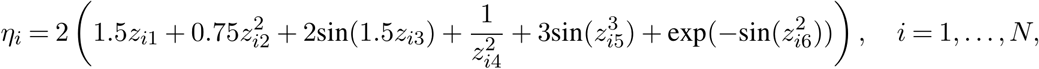

We considered first 6 components of **z**_*i*_’s as significant.

### Scenario 2

Instead of simulating gene expression as independent Gaussian variables, we first generated *N* latent Gaussian vectors **v**_*i*_ ∈ ℝ^*p*^ from a *p* × *p* correlation matrix (Figure 7C). The correlation structure introduces moderately high positive correlations among the first *p/*3 features and negative correlations among the last *p/*3 features, consistent with empirical correlation patterns observed in real scRNA-seq datasets (Figures 4D, 6D). Several established scRNA-seq simulation frameworks, including Splatter (Zappia, Phipson and Oshlack, 2017), SERGIO (Dibaeinia and Sinha, 2020), and scDesign3 (Song et al., 2024) provide end-to-end pipelines for generating realistic scRNA-seq data with important empirical properties such as sparsity and batch effects. However, to our best understanding, these frameworks generally follow a top-down generative structure, fixing cell-type labels a priori and subsequently simulating gene expression conditional on those labels. As a result, they do not define a ground truth for cell-type-predictive gene identification, as discussed in Supplementary Note 1. Nevertheless, these frameworks serve as a valuable reference for benchmarking the empirical properties of externally simulated data. Accordingly, our simulation setting yields an average sparsity of approximately 31%, consistent with sparsity levels observed in scDesign3-simulated data (Song et al., 2024). Furthermore, the imposed correlation structure reflects realistic co-expression patterns characteristic of scRNA-seq data, as illustrated in Figure 7C.

Next, we sampled the cell-specific gene expressions **z**_*i*_ from negative binomial (NB) distributions, *z*_*jm*_ ∼ NB(*ψ*_*jm*_, *ϕ*_*jm*_ = 1), with *ψ*_*jm*_ = exp(1 + *v*_*jm*_). Using the log-transformed and scaled expression matrix 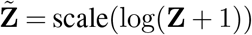, we subsequently considered two designs to generate the outcomes.

### Design 3.

We simulated the outcome logits ***η*** using the ARD kernel equation (3.1). The effect vector **r** had exactly four non-zero entries, with *r*_*m*_ = 1 for *m* ∈ {1, 3, 3*p/*5 + 1, 3*p/*5 + 5} and *r*_*m*_ = 0 otherwise. We fixed 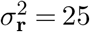 and *p* = 300.

### Design 4.

Finally, we considered another additive model with four significant covariates, but with nonlinear interaction terms,

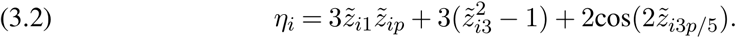

We fixed *N* at 200 and varied *p* between 50, 100 and 400. In particular, setting *p* = 400 creates a popular *N < p* scenario observed in high dimensional data.

For all four designs and corresponding hyperparameter setup, we generated *y*_*i*_’s from a Bernoulli distribution: *y*_*i*_∼ *Bernoulli*(*µ*_*i*_), 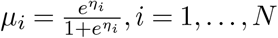. Crucially, choosing the true predictive genes in Designs 1–4 provides us with a practical common ground for simulating all the datasets.

Except for Design 3, we generated 50 replicated datasets per simulation setting. Due to the large sample sizes in Design 3, the number of replicates was reduced to 20 to limit computational cost. To benchmark CBKMR, we considered GBKMR and four per-gene marker identification frameworks – the Wilcoxon rank-sum test, the two-sample *t*-test, Presto (Korsunsky et al., 2019) and DESeq2 (Love, Huber and Anders, 2014b) – as well as two ensemble learning methods: random forest (RF) and XGBoost (XGB), both of which have recently been applied to cell-type classification tasks (Pullin and McCarthy, 2024; Xu et al., 2023). We evaluated marker recovery performance using the area under the precision-recall curve (AUPRC), computed via `average_precision_score` from *scikit-learn* and averaged over replicated datasets (Figure 2A, C, E, G). For CBKMR and GBKMR, we computed computed AUPRC using PIPs as the ranking score. For the per-gene testing frameworks, adjusted *p*-values were used. Since RF and XGB do not yield direct inclusion probabilities, we utilized their gene-specific variable importance scores to computer AUPRC. These methods differ in how gene importance is defined, and their AUPRC values are therefore not always directly comparable. To facilitate a method-agnostic visual comparison of gene-level importance across methods, we further computed standardized importance scores for each method (Figure 2B, D, F, H). In addition to using PIP values for CBKMR and GBKMR, we divided importance scores for RF and XGB by their respective maximum values. For univariate differential expression tests, we used the proportion of replicates in which each gene was declared significant at the 95% confidence level as the standardized importance. Such construction ensures that all standardized importance values lie in [0, 1], providing a common ground for visual comparison.

**Fig 1:**
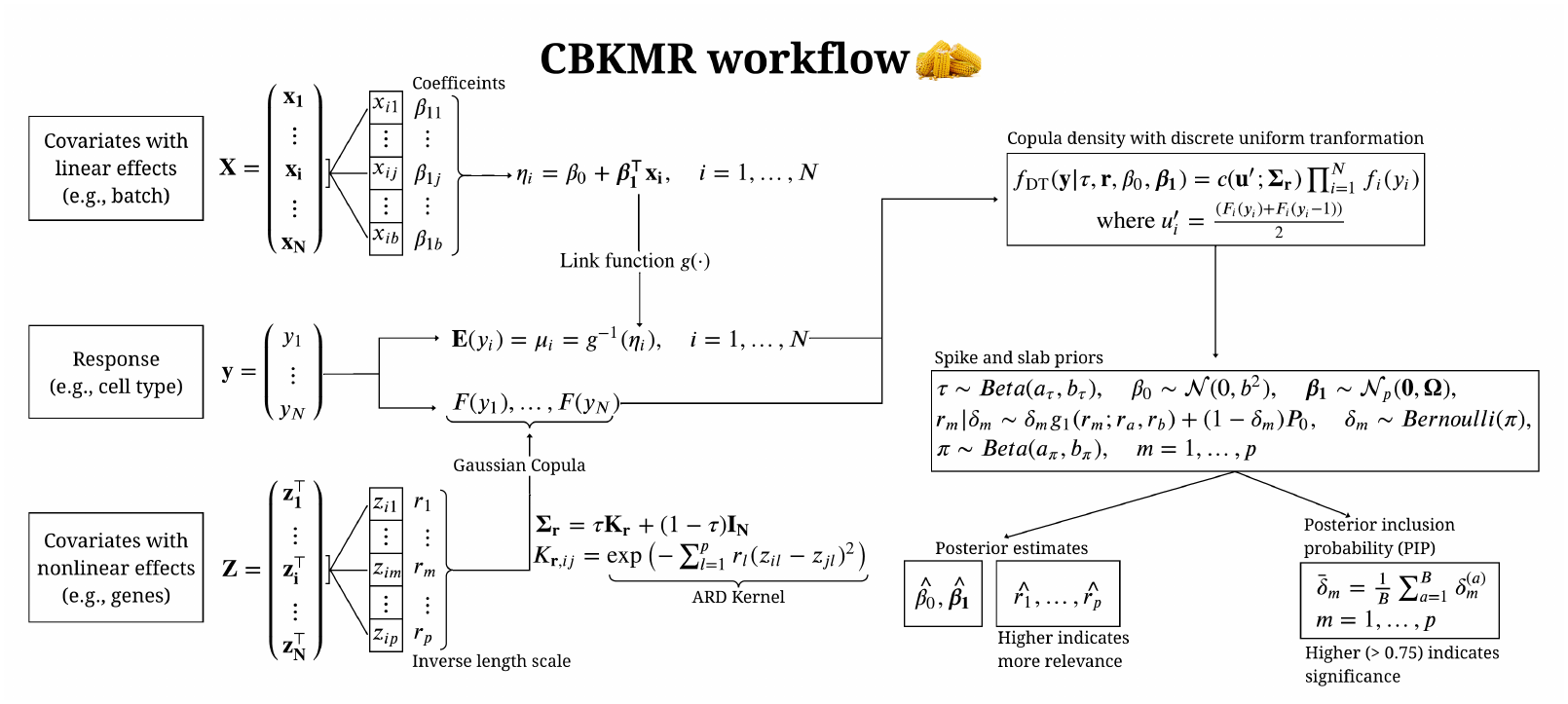
CBKMR workflow diagram with key modeling steps.

**Fig 2:**
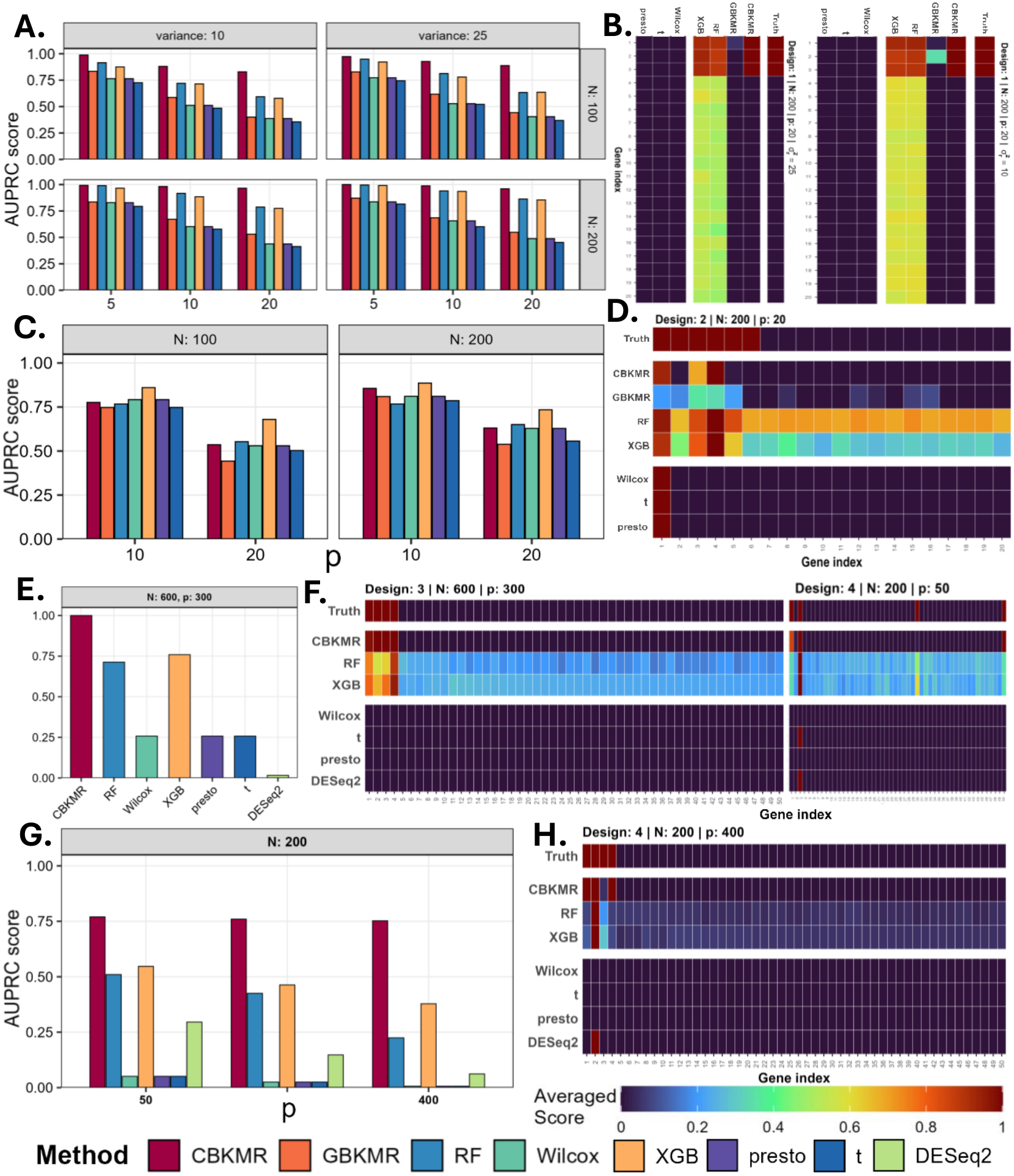
AUPRC and feature importance score comparisons for different designs in Section 3. **A, B**. Selection AUPRC and average importance score corresponding to Design 1. **C, D**. Selection AUPRC and average importance score corresponding to Design 2. **E**. Selection AUPRC corresponding to Design 3. **F**. Average importance score corresponding to two settings for Design 3 and 4 respectively. **G, H**. Selection AUPRC corresponding to *N* = 200 and average importance score corresponding to *N* = 200, *p* = 400. Since Figures **A**–**D** correspond to Designs 1 and 2 where Gaussian distributions were used for gene simulations, they do not contain results for DESeq2.

Figure 2 presents AUPRC scores across all simulation designs. CBKMR consistently out-performed competing methods, in some instances achieving multi-fold improvements over GBKMR (e.g., Figure 2A: 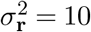. Notably, CBKMR substantially out-performed GBKMR in Design 1 (Figure 2A), despite outcomes being simulated directly from a logistic GBKMR model with an ARD kernel. This aligns with findings by Mad-sen (2009) and Hughes (2015), demonstrating that modeling dependence solely through the marginal link function is often inadequate for discrete outcomes. CBKMR consistently produced higher standardized scores for true variables compared to all univariate DE tests (Figure 2B). The univariate DE tests failed to consistently identify true genes even at high marginal kernel variance 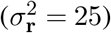, yielding an average score of 0.4 (Figure 2B, Table 3), highlighting their limitation in highly nonlinear settings under small sample sizes.

In Design 2, CBKMR again outperformed all competing methods (Figure 2C). While it assigned nonzero PIP values to some inactive features (Figure 2D), these generally did not exceed 0.2, indicating effective exclusion from the final selected subset. Consistent with these findings, CBKMR recovered more true positive features than univariate DE tests under the complex simulation structure of Design 2 (Table 4). Its performance in Designs 3 and 4 further underscores the robustness of CBKMR in high-dimensional settings. In Design 3, CBKMR achieved near-perfect performance (AUPRC = 1), recovering all four true features (Figure 2E, F, Table 4). By contrast, univariate DE tests such as DESeq2 produced exclusively false positives, highlighting their limitation in high-dimensional settings.

In Design 4, characterized by highly nonlinear gene-outcome relationships and gene-gene interactions, CBKMR consistently achieved higher AUPRC than univariate methods (Figure 2G). CBKMR recovered an average of 2.45 true features even at *p* = 400, whereas DE-Seq2 recovered approximately one true feature on average, incurring a substantially higher false positive rate (Table 4). Naturally, DESeq2 assigned considerably lower importance scores to true features (Figure 2H).

Importantly, CBKMR outperformed RF and XGB in all settings except Design 2, where both ensemble methods achieved higher AUPRC. This difference is likely attributable to how each method distributes importance scores: CBKMR assigns exact zero PIPs to many inactive variables, whereas RF and XGB assign small but nonzero importance scores to all variables by construction. Since AUPRC is sensitive to the full ranking of variables, methods that assign nonzero scores broadly can achieve higher recall at lower precision thresholds, artificially inflating their AUPRC relative to methods that produce sparse, hard-zero assignments. We therefore examined average standardized importance scores to assess whether this AUPRC advantage translates into parsimonious marker selection. Both RF and XGB assigned substantially higher importance scores to inactive features (Figure 2D), limiting their utility for sparse biomarker identification. With increasing *p*, RF and XGB assigned progressively lower importance to inactive features, yet unlike CBKMR, GBKMR, and univariate DE tests, they did not provide a uniform criterion for feature selection thresholding. Given the lower importance scores assigned to inactive features in Figure 2H, one might consider applying a median or upper 75th percentile threshold for marker selection from these models. However, as shown in Section 4.3, in real scRNA-seq datasets where correlation-based filtering retains several redundant markers, RF and XGB assign high importance scores to redundant genes, failing to yield a parsimonious set of predictive marker genes (Table 6a).

Finally, to assess predictive performance, simulated data were partitioned into 80% training and 20% test sets. CBKMR, RF, and XGB were each trained on the training partition and evaluated on the held-out test set, with this process repeated across 20 independent splits of 20 simulated datasets. Since CBKMR is computationally intensive at large sample sizes, posterior samples from a single training split were reused to generate predictions across all 20 test splits within a single replicate; RF and XGB were trained independently on each split. CBKMR achieved an AUPRC of 0.94, comparable to RF (0.99) and XGB (0.98). The marginally higher specificity of RF and XGB was expected, as these models are primarily optimized for prediction rather than variable selection. Taken together, the strong marker selection performance of CBKMR, combined with competitive predictive accuracy, establishes it as an effective tool for sparse biomarker detection in scRNA-seq data.

### 3.1. Efficiency of NNCBKMR in comparison to CBKMR

To evaluate the accuracy and computational efficiency of NNCBKMR on large datasets, we considered ten simulation settings: six combinations under Design 3, with *N* ∈ {1000, 2000} and *p* ∈ {50, 100, 150}, and four (*N, p*) combinations under Design 2, with *N* ∈ {700, 1000} and *p* ∈ {10, 20}. We avoid running GBKMR in this setting due to its prohibitive computational cost. For Design 3, we evaluate NNCBKMR for only *k* = 40 neighbors, while for Design 4, we evaluated NNCBKMR for *k* = 5, 10, 20 and 40 neighbors. As shown in Figure 3A, NNCBKMR with *k* = 40 achieved performance comparable to CBKMR across varying dimensions for Design 3, perfectly selecting all 4 of the true positive features (Table 5). In contrast, univariate DE approaches such as Wilcoxon rank-sum test and presto recovered 3 out of the 4 features. While DESeq2 correctly recovered an average of 2 genes, it produced 6 false positives on ane average (Table 5). Naturally, all the univariate DE approaches achieved higher power in the presence of larger sample size that was further reflected in their importance scores (Figure 3B). However due to the presence of the gene-gene interaction in Design 4, their power noticably dropped even for *p* = 20, resulting in lower AUPRC values (Figure 3C) and vanishing gene-specific importance scores (Figure 3D). While DESeq2 still maintained more true positives compared to the other univariate DE tests (Table 5), it consistently produced larger number of false positives. In contrast, NNCBKMR with a nominal neighbor size of *k* = 5 outperformed existing DE methods, achieving almost accurate AUPRC of ∼1 (Figure 3C). Under the presence of larger sample sizes, RF and XGB showed consistent performance, assigning high importance to relevant genes for both designs. However, similar to our earlier finding, these models could not produce a uniform cutoff criteria for producing parsimonious marker genes. Finally, both NNCBKMR and CBKMR achieved a high prediction accuracy with an AUPRC value of 0.91, consistent with their RF and XGB counterparts, who produced AUPRC of 0.98 and 0.97 respectively.

**Fig 3:**
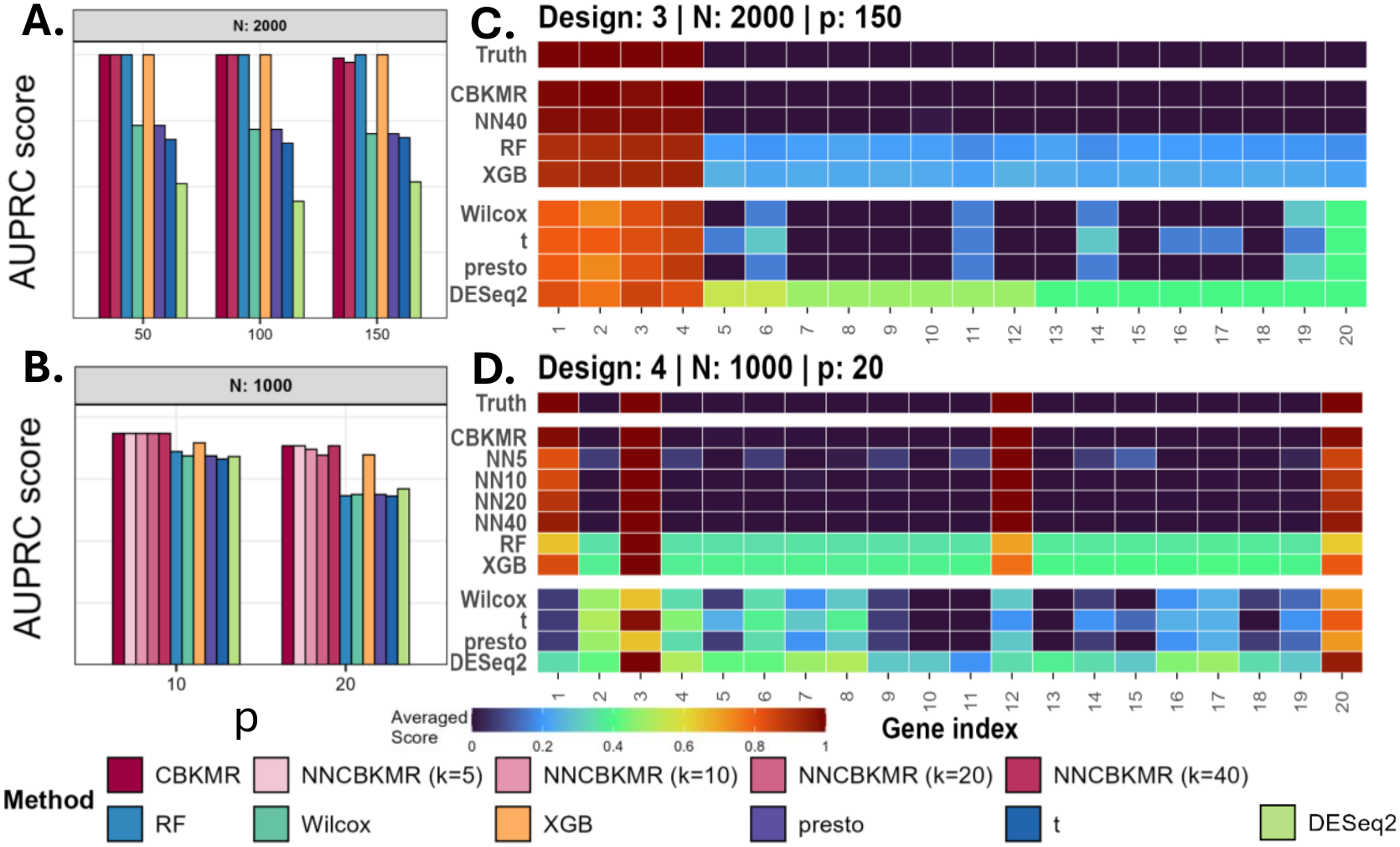
Comparison between CBKMR and NNCBKMR using precision recall curves. **A**. AUPRC scores at *N* = 2000 for Design 3. **B**. Importance scores corresponding to **A. C**. AUPRC scores for *N* = 1000 at Design 4. **D**. Importance score corresponding to **C**. For **B** and **D** NNCBKMR has been alised as NN.

Furthermore, even at *N* = 2000 and *p* = 150 under Design 4, NNCBKMR correctly identifies an average of 3.95 out of 4 true markers, demonstrating robust marker recovery in high-dimensional settings. While conventional per gene tests such as Wilcoxon rank-sum tests and DESeq2 achieve high specificity as *N* increases, their performance suffer from the nonlinear simulation structure of Design 4 (Figure 3B), which underscores their limitation in identifying nonlinear gene-outcome interaction even under large sample sizes. Additionally, NNCBKMR also offers notable computational advantage over CBKMR. As reported in Table 1, NNCBKMR with *k* = 20 achieveed approximately 1.5-fold runtime reduction relative to CBKMR at *N* = 700. At *k* = 40, the modest overhead of NNCBKMR relative to CBKMR at moderately large sample sizes is attributable to its manually parallelized computational routine, which does not yet match the speed of the optimized matrix operations implemented via the `Armadillo C++` library. However, as *N* increases, NNCBKMR (*k* = 40) rapidly surpassed standard CBKMR in speed, underscoring its practical utility for large samples. For instance, at *p* = 150, NNCBKMR achieved approximately 1.3-fold and 2.4-fold runtime reductions relative to CBKMR at *N* = 1000 and *N* = 2000, respectively. As noted in Section 2.5, the runtime of NNCBKMR scales polynominally with *p*. At *N* = 2000, increasing *p* from 50 to 150 resulted in approximately 1.8-fold and 2.3-fold runtime increases for CBKMR and NNCBKMR, respectively, reflecting a marginally greater sensitivity of NNCBKMR to feature dimensionality. Nevertheless, following the recommendation of sure independence screening in practical high-dimensional settings (Fan and Lv, 2008) we can reduce the feature space to approximately 150–200 candidate genes, maintaining NNCBKMR’s feasibility under moderate values of *p*. The peak memory consumption of both CBKMR and NNCBKMR remains in the order of megabytes. For *N* = 2000, the memory usage for CBKMR roughly doubles with respect to NNCBKMR. This is likely attributable to the complete matrix calculation performed by CBKMR. NNCBKMR, through its construction of approximate density function avoids such large matrix computations. Besides its marker selection capability, NNCBKMR also achieves predictive performance comparable to CBKMR across both designs (Figure 3C). Collectively, these results validate NNCBKMR as a computationally efficient and statistically robust approximation of CBKMR for moderately large feature dimensions.

**TABLE 1.** Completion time (T, in minutes) and peak memory usage (PM, in megabytes) comparison between CBKMR and NNCBKMR, averaged over 20 replicated simulations. GBKMR took 128 minutes (2 hours) to complete 2,000 iterations for N = 700 and p = 10. We did not track peak memory usage for GBKMR, since it approximately takes 5000 iterations to converge, which is a prohibitive task for even the moderately large dataset.

| N | p | NNCBKMR |  |  |  |  |  |  |  | CBKMR |  |
|---|---|---|---|---|---|---|---|---|---|---|---|
|  |  | k = 5 |  | k = 10 |  | k = 20 |  | k = 40 |  |  |  |
|  |  | T | PM | T | PM | T | PM | T | PM | T | PM |
| 700 | 10 | 3.46 | 190 | 3.8 | 211 | 4.47 | 185 | 8.64 | 206 | 5.93 | 188 |
|  | 20 | 7.23 | 216 | 7.9 | 211 | 10 | 201 | 17.9 | 212 | 12.7 | 208 |
| 1000 | 10 | 4.65 | 201 | 5.2 | 210 | 6.28 | 207 | 11.7 | 206 | 15.7 | 232 |
|  | 20 | 10.1 | 201 | 11 | 225 | 13.7 | 214 | 26.4 | 209 | 32.7 | 232 |
|  | 50 | – |  |  |  |  |  | 30.3 | 212 | 45.3 | 263 |
|  | 100 |  |  |  |  |  |  | 64.7 | 228 | 94.3 | 270 |
|  | 150 |  |  |  |  |  |  | 68.6 | 240 | 92.8 | 280 |
| 2000 | 50 | – |  |  |  |  |  | 81.6 | 248 | 244.2 | 496 |
|  | 100 |  |  |  |  |  |  | 132.8 | 249 | 351.7 | 533 |
|  | 150 |  |  |  |  |  |  | 191.3 | 255 | 441.1 | 560 |

## 4. Real data analysis

In this section, we validate the utility of CBKMR on three scRNA-seq datasets. These datasets commonly contain redundant features alongside mito-chondrial and ribosomal RNA transcripts, necessitating careful preprocessing prior to statistical analysis. Unless otherwise stated, we perform the following preprocessing steps uniformly on all of the three datasets. All the data preprocessing has been performed with *R* packages such as Seurat, scran and scuttle.

### (1) Cellular quality control (QC)

After removing all the mitochondrial and ribosomal RNAs, we performed QC of the datasets using `perFeatureQCMetrics` and retained the genes passing a threshold. For the first two datasets, we used a threshold value of 0.01, while for the pancreatic data in Section 4.3, we used a threshold of 0.001 due to its higher sparsity from droplet-based sequencing.

### (2) Log-normalization and variance stabilization

Next, we performed log-normalization under a scale factor of 10000 in the `NormalizeData` from *Seurat*. Subsequently, we performed variance stabilization using the `FindVariableFeatures` from *Seurat* and kept the top 2000 variable features.

### (3) FDR control and correlation filtering

Next, for each one-versus-rest cell-type comparison, we performed univariate DE analyses using the Wilcoxon rank-sum test implemented in *scran*, and retained genes at an FDR threshold of 0.05. In high-dimensional settings, many genes are either redundant or weakly associated with cell-type identity; FDR-based prescreening reduces the feature space to a manageable set of candidate markers prior to kernel-based modeling. In Section 4.3, this threshold was relaxed to 0.99 (retaining nearly all genes) to evaluate CBKMR under a larger and less filtered feature set. Finally, we applied a correlation filtering using the `findCorrelation` function (Kuhn, 2008) and removed genes with pairwise correlation exceeding 0.75, as high collinearity among covariates can destabilize the estimation of inverse length-scale parameters in the ARD kernel. In Sections 4.2 and 4.3, we further reduced this threshold to 0.5 to assess the sensitivity of CBKMR to correlated covariates. In the Supplementary Note, we provide a discussion on the importance of this correlation filtering step, a widely adopted preprocessing approach for reducing multicollinearity and improving numerical stability of kernel-based models.

In addition to data preprocessing, we also applied a post-processing on the posterior samples of CBKMR and analysed the results based on the selected genes (i.e., PIP > 0.9) with posterior mean of *r*_*m*_ *>* 0.1.

### 4.1. Analysis of Type 2 diabetes islet-cell data

Lawlor et al. (2017) generated single-cell transcriptomes (26,616 genes) from 638 human pancreatic islet cells collected from both non-diabetic and type 2 diabetic (T2D) donors. Their analysis identified distinct transcriptional signatures for the major endocrine islet cell types (e.g., alpha, beta, delta, and gamma/PP cells) and revealed rare cellular states associated with islet dysfunction. Importantly, the dataset highlighted cell-type–specific expression changes in T2D that were not detectable in bulk islet profiling. In their downstream analyses, the authors relied on a curated set of canonical marker genes to annotate cell types, including *INS* (beta), *GCG* (alpha), *SST* (delta), *PPY* (gamma/PP), *PRSS1* (acinar), *COL1A1* (stellate), and *KRT19* (ductal). Af-ter minimal quality-control filtering, we retained 604 cells with the following highly skewed distribution of cell-type labels: alpha (238), beta (252), acinar (23), delta (20), ductal (23), gamma/PP (17), stellate (19), and other (12). Notably, *SST* and *PPY* were absent from the filtered dataset after removing highly correlated genes. To retain scientific validity of the results, we added the missing canonical markers to the filtered dataset, resulting in a final set of 132 genes in the feature expression matrix. We evaluated CBKMR by examining whether it could successfully recover the canonical marker for every cell type.

CBKMR identified 13 genes with PIPs exceeding 0.95 and estimated inverse lengthscales *r*_*m*_ *>* 0.1. In Figs. 4A and 4B, we show the mean expression of all genes within each cell type and highlight the 13 selected genes. Notably, CBKMR identified six of seven canonical markers as the top-ranked feature across six cell types, with the exception of *COL1A1* in stellate cells. As shown in Fig. 4C, the genes form pronounced clusters, likely due to the relatively permissive correlation threshold of 0.75 used during preprocessing. Figure 4D further shows that most genes in the highlighted *COL1A1* branch are strongly correlated with *COL1A1*. In particular, *WNT5A* and *COL5A2*, both selected by CBKMR for stellate cells, are substantially correlated with *COL1A1* (approximately 0.6) and have small inverse lengthscales (0.3 and 0.224, respectively; Fig. 4E). This pattern indicates that CBKMR may occasionally select one or more representative genes from a correlated block, an effect that may be further amplified by class imbalance (19 stellate cells vs. 585 other cells). CBKMR’s strong overall performance underscores its utility for complex real datasets in which genes exhibit substantial correlation. While the correlation threshold could be tightened to yield a smaller candidate gene set, we intentionally used this more extreme setting to evaluate performance in a highly correlated framework. For random forests and XGBoost, feature selection based on variable importance is inherently ad hoc; therefore, we focus on the top-ranked gene identified by each method. XGBoost selected the correct marker as the top gene for every cell type. Random forests selected the correct marker for every cell type except stellate cells, for which it flagged *VCAN*. However, results from both methods varied across random seeds and occasionally failed to identify some marker genes, underscoring the need for caution when relying on machine learning–based feature rankings to develop predictive marker panels, such as those implemented in recent software packages (Nelson, Riva and Cvejic (2022)).

**Fig 4:**
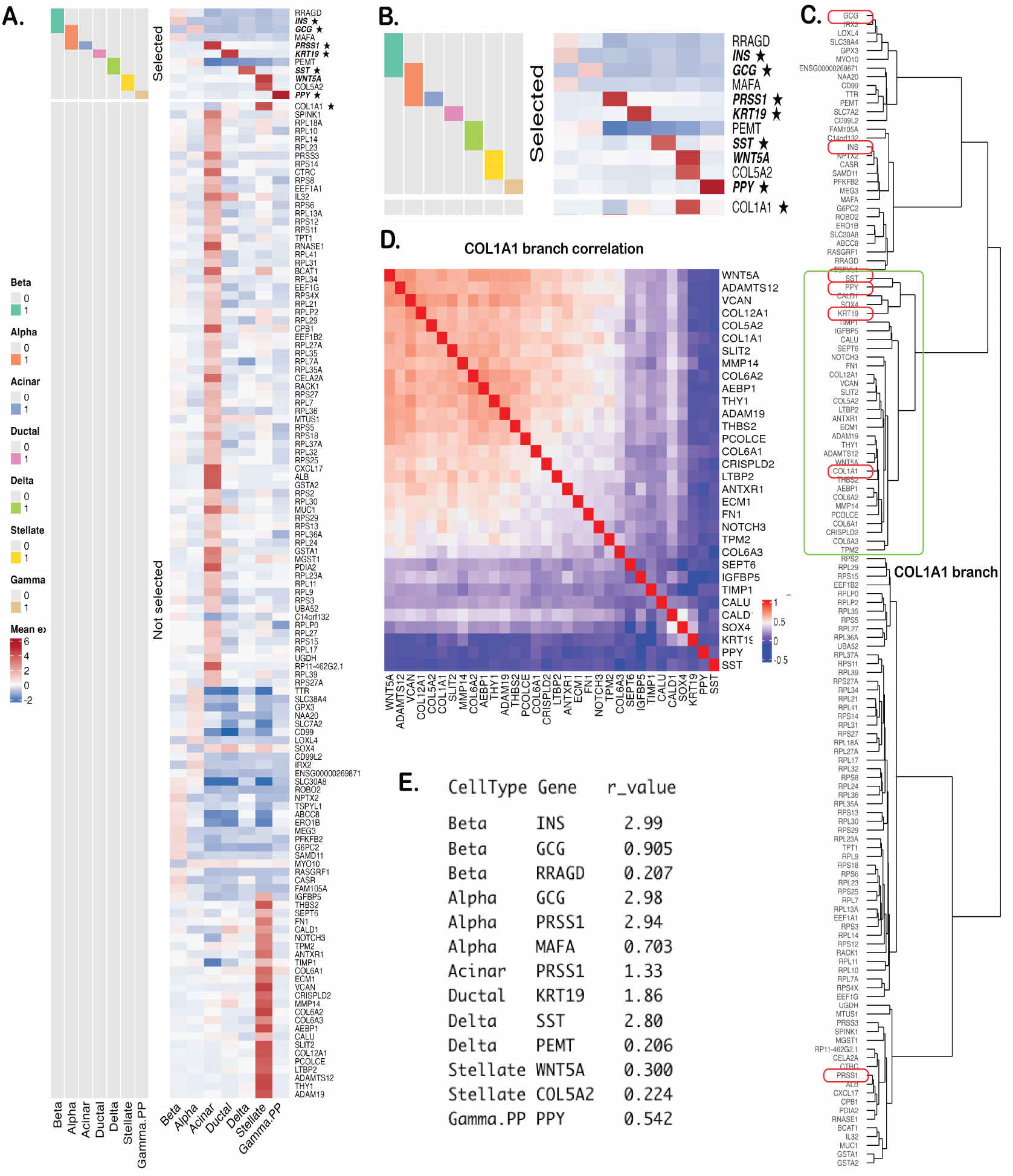
**A**. Heatmap of the average expression of 132 genes across seven cell types. Pathologist-annotated markers for each cell type are indicated with an asterisk (*). CBKMR-selected genes are shown for each cell type (0 = not selected, 1 = selected), with the *top* feature high-lighted in bold. **B**. Zoomed view of the CBKMR-selected genes. **C**. Hierarchical clustering of genes; red boxes denote the markers, and the green box highlights the branch containing *COL1A1*. **D**. Correlation heatmap of genes within the *COL1A1* branch. **E**. Estimated inverse lengthscales (relevance) for the selected genes.

### 4.2. Analysis of Ncx1 ^−/−^ and wild type mouse embryos

We analyzed a publicly available dataon mouse embroys generated by Azzoni et al. (2021) to study activation trajectories at the onset of circulation. Circulation refers to the heart-driven pumping of blood, which generates mechanical forces in the embryo that are essential for triggering critical developmental processes, such as the Endothelial-to-Hematopoietic Transition (EHT). In particular, EHT is a foundational developmental process essential for life, as it is the original source of all lifelong blood and immune cells. During EHT, specialized endothelial (vessel-lining) cells transform, detach, and are ultimately released into the body as the first hematopoietic stem cells. Wild-type embryos possess a normal heartbeat and hemodynamic flow, allowing EHT to be activated through proper circulation. Ncx1^−*/*−^ embryos, on the other hand, do not go through a proper circulation process, which in turn leads to impaired EHT activation. Therefore, the comparison between Ncx1^−*/*−^ and wild-type embryos is critical for identifying the genes and functional processes that are crucial for circulation.

We obtained the gene expression data and cell type annotations from the European Bioin-formatics Institute single cell expression atlas website EMBL-EBI (2025). For our analysis, we only kept the Ncx1^−*/*−^ and wild-type cells. The final dataset contained *N* = 444 observations and 353 genes with 244 Ncx1^−*/*−^ and 200 wild-type cells. CBKMR produced PIP values of 1 for 8 out of the 50 genes, rendering those genes significant for cell type prediction. While this dataset presents a significantly large number of DE genes (368, Figure 5B), the performance of CBKMR in identifying a small feature set highlights its utility in high dimensional data. Since this dataset does not represent any canonical marker, we investigated scientific validity of the selected genes. Among the selected genes in the final dataset, *Cd44* was previously reported as an essential marker for EHT (Oatley et al. (2020)), consistent with the authors’ finding that *Cd44* is generally downregulated in the case of impaired EHT. Hierarchical clustering showed that CBKMR picked Lcp2, a feature closely clustered with Cd44 (Figure 5A). Notably, Lcp2 is an established regulator of immunoreceptor signaling, a key component of EHT (Iyer et al., 2022; Michaels, Durland and Zandstra, 2023). Additionally, Sox17 has been noted as a key component in EHT, and is downregulated on successful hematopoitic transition. CBKMR picks Dusp6, a gene appearing in the clustered block of Sox17 (Figure 5A). CBKMR also picks Dll4, a gene that behaves similar to Sox17 during EHT. Among other selected genes, *Sox9* was closely related to a key EHT transcription factor *Runx1* in other studies (Shah et al. (2025)). We further increased the correlation cutoff to 0.5, resulting in 289 genes. Even under a stricter threshold, CBKMR picked Ankrd37, a gene adjacent to Cd44, validating its robustness under varied correlation cutoff. Importantly, while *Sox9* was selected in both correlation cutoffs, posterior mean of *r*_*m*_ was 0.36 (Figure 5B) in the correlation thresholding of 0.75, slightly lower than the posterior mean of 0.52 at correlation thresholding of 0.5 (Figure 5B). We also performed random forests and XGBoost on the filtered data. Random forest assigned highest importance score to Dll4, while XGBoost assigned third highest importance to it. Notably, however utilizing the *µ* + 3*σ* cuttof discussed in Xu et al. (2023) retained only eight genes in random forest and 5 genes in XGBoost, none of which selected the other important genes related to Sox17 and Cd44, highlighting the limitation of these methods in identifying minimal predictive feature sets. Furthermore, to test the accuracy of CBKMR, we made a 90-10 split of the dataset and evaluated the prediction accuracy of CBKMR in test dataset. Similar to its simulation counterparts, we only trained CBKMR on one training split and reused them in the other. As shown in Table 2, CBKMR achieved suitable AUPRC compared to the prediction-based random forest and XGBoost models.

**TABLE 2.** Prediction AUPRC for mouse embryo (Dataset 2) and pancreatic cell (Dataset 3) data.

| Method | Dataset 2 | Dataset 3 |
|---|---|---|
| CBKMR | 0.983 | 0.913 |
| RF | 994 | 0.982 |
| XGBoost | 0.994 | 0.976 |

**TABLE 3.**
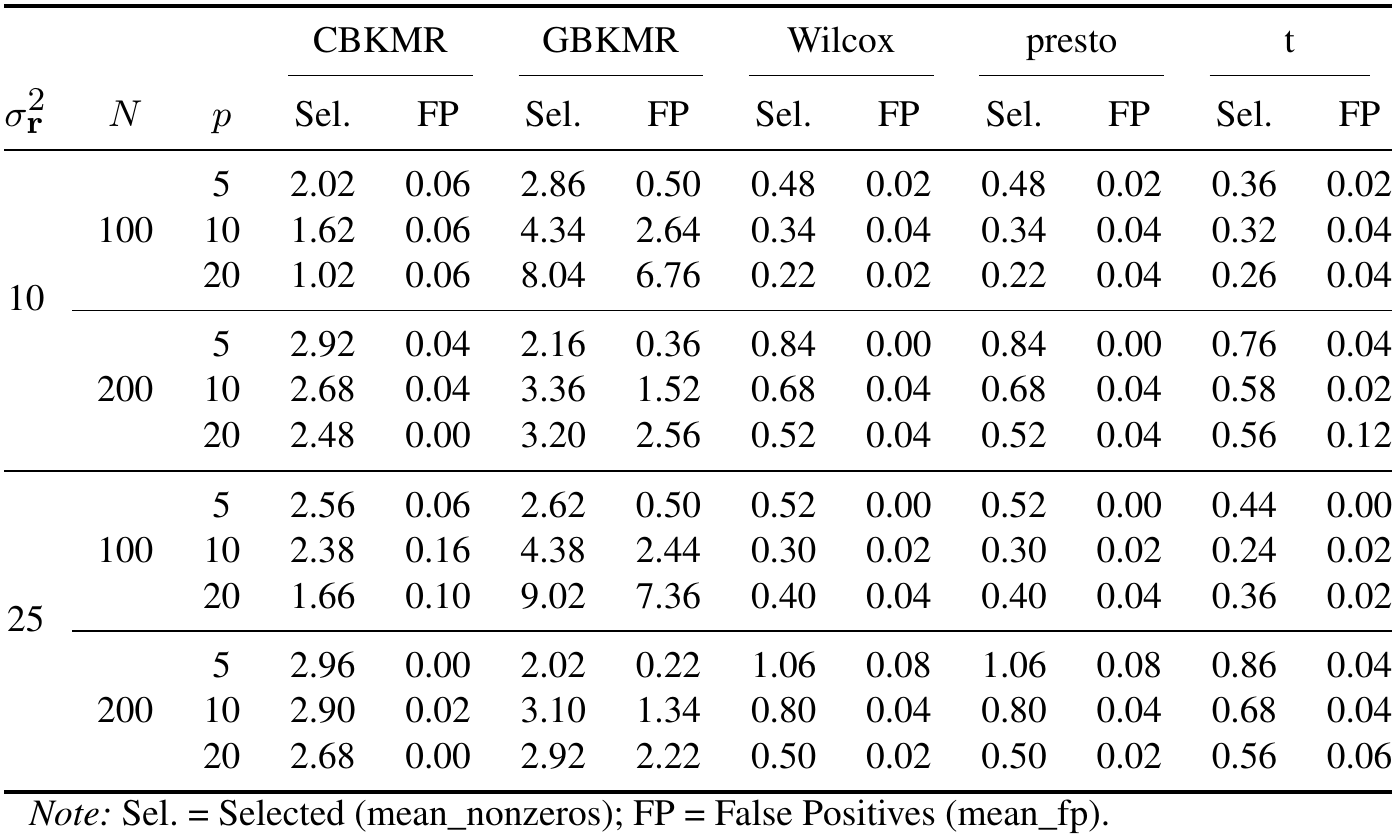
Average Number of selected features and false positives by method for Design 1.

**TABLE 4.** Average number of selected features and false positives by method across designs, sample sizes, and feature dimensions.

|  |  |  | CBKMR |  | GBKMR |  | Wilcox |  | presto |  | t |  | DESeq2 |  |
|---|---|---|---|---|---|---|---|---|---|---|---|---|---|---|
| Design | *N* | *p* | Sel. | FP | Sel. | FP | Sel. | FP | Sel. | FP | Sel. | FP | Sel. | FP |
| 2 | 100 | 10 | 2.5 | 0 | 4 | 1 | 1 | 0 | 1 | 0 | 1 | 0 | – | – |
|  |  | 20 | 2 | 0 | 7.5 | 5 | 0 | 0 | 0 | 0 | 0 | 0 | – | – |
|  | 200 | 10 | 3 | 0 | 2 | 1 | 2 | 0 | 2 | 0 | 1 | 0 | – | – |
|  |  | 20 | 2 | 0 | 2 | 0.5 | 1 | 0 | 1 | 0 | 1 | 0 | – | – |
| 3 | 600 | 300 | 3.5 | 0 | – | – | 1.0 | 0 | 1.0 | 0 | 1.0 | 0 | 3.0 | 3 |
|  | 700 | 400 | 4.0 | 0 | – | – | 2.0 | 0 | 2.0 | 0 | 1.5 | 0 | 6.5 | 5 |
| 4 | 200 | 50 | 2.55 | 0 | – | – | 0.10 | 0.05 | 0.10 | 0.05 | 0.25 | 0.10 | 2.40 | 1.30 |
|  |  | 100 | 2.45 | 0 | – | – | 0.05 | 0.05 | 0.05 | 0.05 | 0.15 | 0.05 | 2.90 | 1.75 |
|  |  | 400 | 2.40 | 0 | – | – | 0.05 | 0.05 | 0.05 | 0.05 | 0.05 | 0.05 | 5.25 | 4.25 |
|  | 400 | 50 | 2.90 | 0 | – | – | 0.30 | 0.15 | 0.30 | 0.15 | 0.80 | 0.30 | 3.20 | 1.85 |
|  |  | 100 | 3.15 | 0 | – | – | 0.15 | 0.10 | 0.15 | 0.10 | 0.70 | 0.40 | 4.35 | 3.20 |

*Note:* Sel. = mean number of selected features; FP = mean number of false positives.

**TABLE 5:**
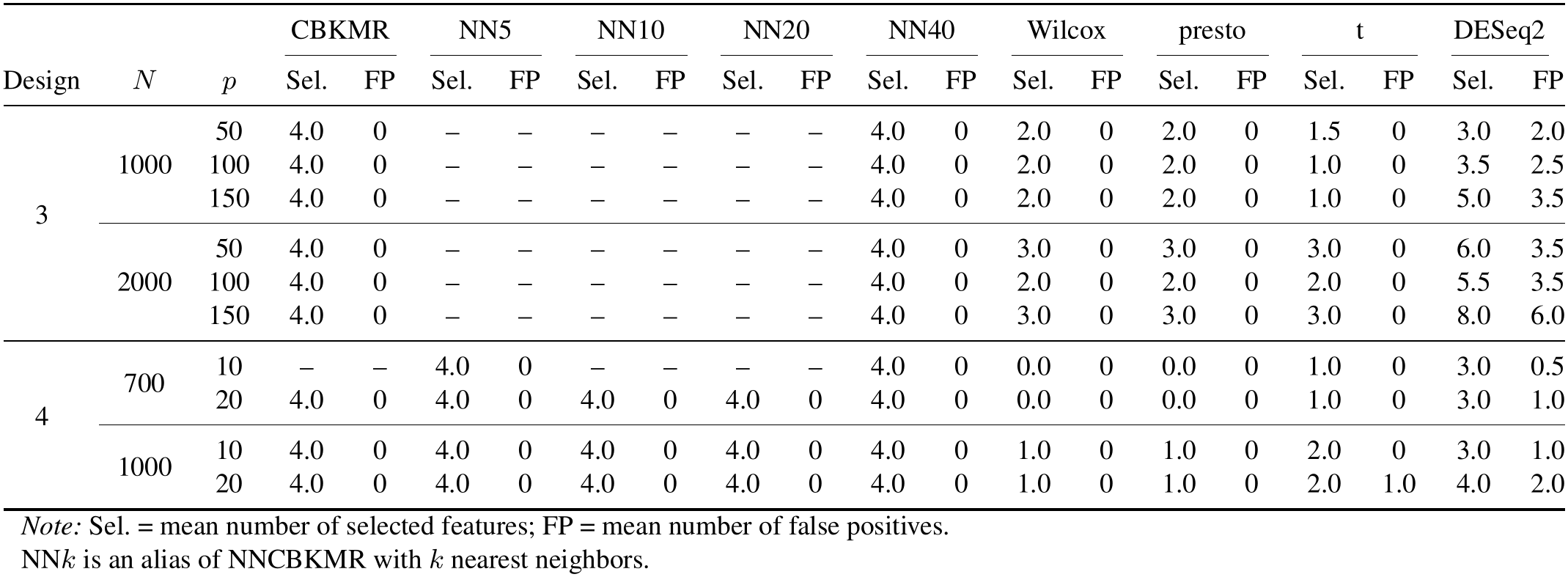
Average number of selected features and false positives by method for NNCBKMR.

**TABLE 6.** Top 10 features selected by feature importance for beta, alpha, and acinar cells in Baron’s pancreatic cell dataset.

| (a) Random Forest |  |  |  |  |  | (b) XGBoost |  |  |  |  |  |
|---|---|---|---|---|---|---|---|---|---|---|---|
| beta |  | alpha |  | acinar |  | beta |  | alpha |  | acinar |  |
| Gene | Imp. | Gene | Imp. | Gene | Imp. | Gene | Imp. | Gene | Imp. | Gene | Imp. |
| INS | 39.22 | GCG | 332.02 | CPA1 | 14.87 | INS | 0.78 | GCG | 0.72 | CPA1 | 0.55 |
| ADCYAP1 | 26.40 | CLU | 102.20 | REG1P | 10.27 | IAPP | 0.05 | CLU | 0.11 | MGST1 | 0.11 |
| IAPP | 23.97 | IRX2 | 69.54 | RNASE1 | 9.82 | ADCYAP1 | 0.04 | IRX2 | 0.03 | RNASE1 | 0.10 |
| HADH | 22.92 | PCSK2 | 14.28 | MGST1 | 9.38 | HADH | 0.02 | LITAF | 0.02 | ALDOB | 0.05 |
| IGF2 | 14.70 | GPX3 | 6.30 | ANPEP | 9.03 | SERPING1 | 0.01 | PPY | 0.01 | ANPEP | 0.04 |
| CLU | 14.28 | LITAF | 5.94 | ALDOB | 9.02 | VIM | 0.01 | CYSTM1 | 0.01 | CD44 | 0.02 |
| HSPB1 | 11.88 | VGF | 5.20 | CD44 | 8.33 | SST | 0.01 | VGF | 0.01 | NBL1 | 0.01 |
| GCG | 11.47 | SLC7A2 | 4.62 | CTRL | 8.30 | CLU | 0.01 | IFITM3 | 0.01 | MUC1 | 0.01 |
| C1QL1 | 10.00 | PPY | 3.19 | UGDH | 8.29 | IFITM3 | 0.01 | SLC7A2 | 0.01 | CD99 | 0.01 |
| ERO1B | 9.90 | CYSTM1 | 2.87 | HLA-A | 7.85 | ZFP36L1 | 0.01 | GPX3 | 0.01 | SDC4 | 0.01 |

*Note for (a):* Following the μ + 3σ rule of Xu et al. (2023), only 6 genes can be retained from acinar cell types (cut-off = 8.81), all of which come from an extremely correlated block of *CPA1* (Figure 6C), resulting in redundant markers and providing little information on genes that are less correlated with *CPA1* but may become predictive of cell type.

**TABLE 7.**
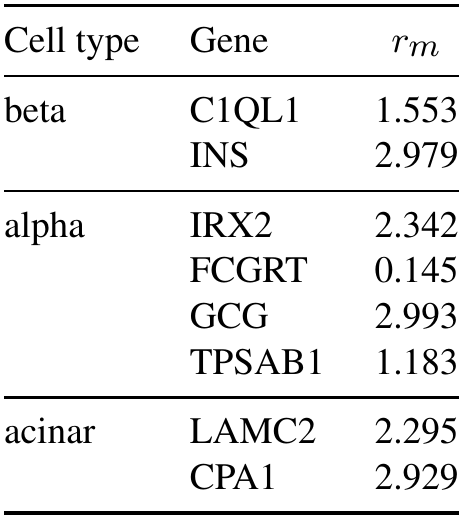
Selected genes using NNCBKMR in Section 4.3 and their rm values by cell type.

| Cell type | Gene | *r*<sub>*m*</sub> |
|---|---|---|
| beta | C1QL1 | 1.553 |
|  | INS | 2.979 |
| alpha | IRX2 | 2.342 |
|  | FCGRT | 0.145 |
|  | GCG | 2.993 |
|  | TPSAB1 | 1.183 |
| acinar | LAMC2 | 2.295 |
|  | CPA1 | 2.929 |

**Fig 5:**
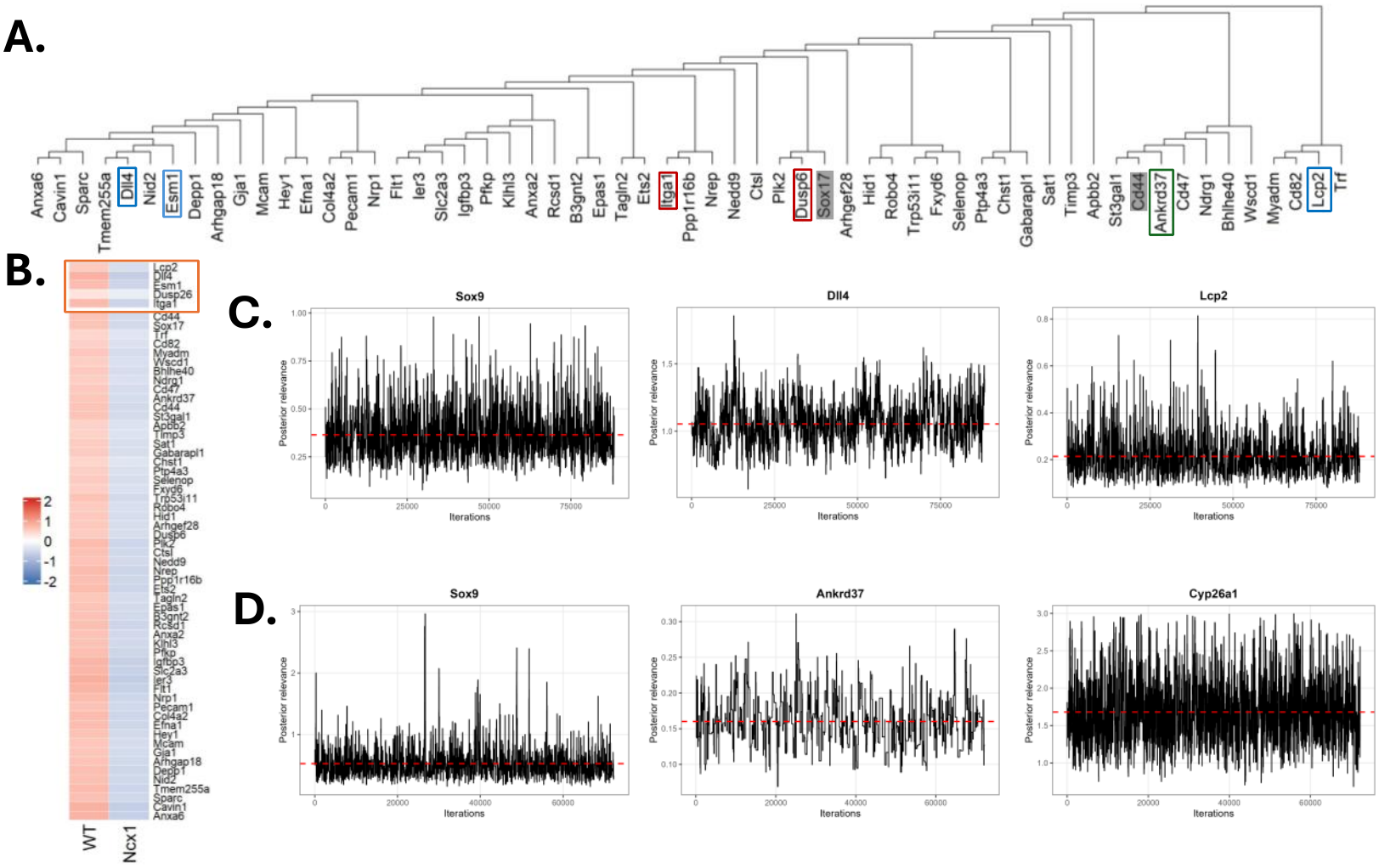
CBKMR results from Ncx1^−*/*−^ mouse embryo data in Section 4.2. **A**. A sub-tree of genetic features containing biologically Cd44, Sox17 that contain selected (colored boxes) and several unselected genes. Genes with red boxes were selected in CBKMR for both 0.5 and 0.75 correlation cutoff, genes with blue and green boxes were selected exclusively at 0.75 and 0.5 cutoff respectively. **B**. Scaled mean expression of the genes in the sub-tree at **A** for both wilde type (WT) and Ncx1^−*/*−^ cells, with red box denoting the selected genes. **C**. Posterior plot of *r*_*m*_ for several relevant genes at correlation cutoff 0.75. **D**. Posterior plot of *r*_*m*_ for several relevant genes at correlation cutoff 0.5.

### 4.3. Analysis of human pancreatic cell types

Single-cell transcriptomic data from four human cadaveric donors (Baron et al. (2016)) were utilized to provide a comprehensive base-line of adult pancreatic cell populations. Data were acquired via the `panc8` dataset (version 3.0.2) using the *SeuratData* R package (Satija, Hoffman and Butler (2026)). The full dataset consists of 34,643 genes measured across 8,569 cells, characterized by an imbalanced distribution dominated by beta, alpha, acinar, and ductal cell types. All the annotations relied on a curated, pathologist-annotated marker gene set, maintaining high consistency with the markers described in Section 4.1. To reduce computational burden, we performed a stratified random sampling approach to extract 1,500 cells, strictly maintaining the original population proportions (yielding a subset containing 439 beta, 404 alpha, and 202 acinar cells). After performing the univariate DE analysis and removing highly correlated genes with correlation cut-off of 0.75, the final dataset contained 161 genes including the canonical markers.

We subsequently applied CBKMR independently to the three most abundant populations (beta, alpha, and acinar cells) using a one-versus-all binary classification framework. CBKMR successfully identified canonical, pathologist-annotated markers for beta (*INS*) and alpha (*GCG*) cells, with posterior mean of *r*_*m*_ close to 3, reflecting importance of these markers (Figure 6A and 6E). Notably, CBKMR did not pick *CPA1*, the canonical marker gene for acinar cell types. Hierarchical clustering showed that among the selected genes by CBKMR, several biologically relevant genes such as *RNASE1* and *CD44* clustered together with *CPA1* with posteior mean of *r*_*m*_ 0.26 and 2.69 respectively (Figure 6B and 6C). Both of these genes showed strong correlation with *CPA1* ( 6D). Therefore, we hypothesized that non-selection of *CPA1* could be due to the identifiability issue arising from the permissive correlation filtering. To further test this claim, we increased the correlation threshold to 0.5, resulting in 118 genes in the final feature matrix. Under the stringent correlation cutoff, CBKMR selected all the canonical markers including *CPA1* (Figure 6F and 6G). In particular, while the posterior mean of *r*_*m*_ was 2.96 for *CPA1* under stringent correlation cut-off of 0.5, the same for its adjacent gene *RNASE1* attained only a posterior mean of 0.25 under the permissive cut-off of 0.75, consistent with identifiability issue observed in Section 5. Among the remaining selected genes from CBKMR under the permissive correlation cut-off, the model identified biologically relevant secondary features. For instance, *TPSAB1*, a known immune cell marker associated with stromal invasion in pancreatic cancer patients (Crnogorac-Jurcevic et al. (2013)) achieved a moderately large mean posterior mean of 1.48 (Figure 6E). While not studied in the context of this specific dataset, *IRX2* has historically shown strong association with alpha cells (Dorrell et al., 2011). Notably, we also applied NNCBKMR to the gene expression matrix. Similar to CBKMR, NNCBKMR was also able to capture all the marker genes with high relevance (Table 7), establishing the accuracy on the approximation framework not only on the simulated dataset, but also on the real scRNA-seq datasets. While both Random Forest and XGBoost correctly assigned the highest importance to the primary established marker genes, their subsequent top rankings were dominated by features highly collinear with these primary markers. This correlation bias is a well-documented limitation of tree-based ensembles, which tend to distribute feature importance across correlated variables rather than isolating independent signals (Toloşi and Lengauer, 2011). For instance, in the acinar cell type classification, the top-ranked genes in both random forest and XGBoost consisted largely of redundant features that were strongly correlated with the primary marker *CPA1* (Table 6a, Figure 6A, C). Consequently, the feature sets identified by these ML models are highly redundant, failing to yield a parsimonious biological signature. In contrast, CBKMR identified a parsimonious set of orthogonal markers that define a clear decision boundary, successfully isolating non-redundant signals even within a highly correlated gene expression space. We further tested the performance of CBKMR on the 20 replicated held out test datasets of size 100, sampled from the remaining 6500 samples not utilized in the training. CBKMR showed reasonable AUPRC in comparison to random forest and XGBoost (Table 2). Finally, to assess the performance of CBKMR under a much larger gene set, we relaxed the FDR to 0.99, and used a correlation cut-off of 0.5 to prevent unwanted curse of dimensionality in large dimensions, which resulted in 378 genes in the final dataset. Even under the high dimensional setup, CBKMR properly selected the three canonical marker genes including *CPA1* (Figure 6H, 8), underscoring its robustness in high dimensional setting.

**Fig 6:**
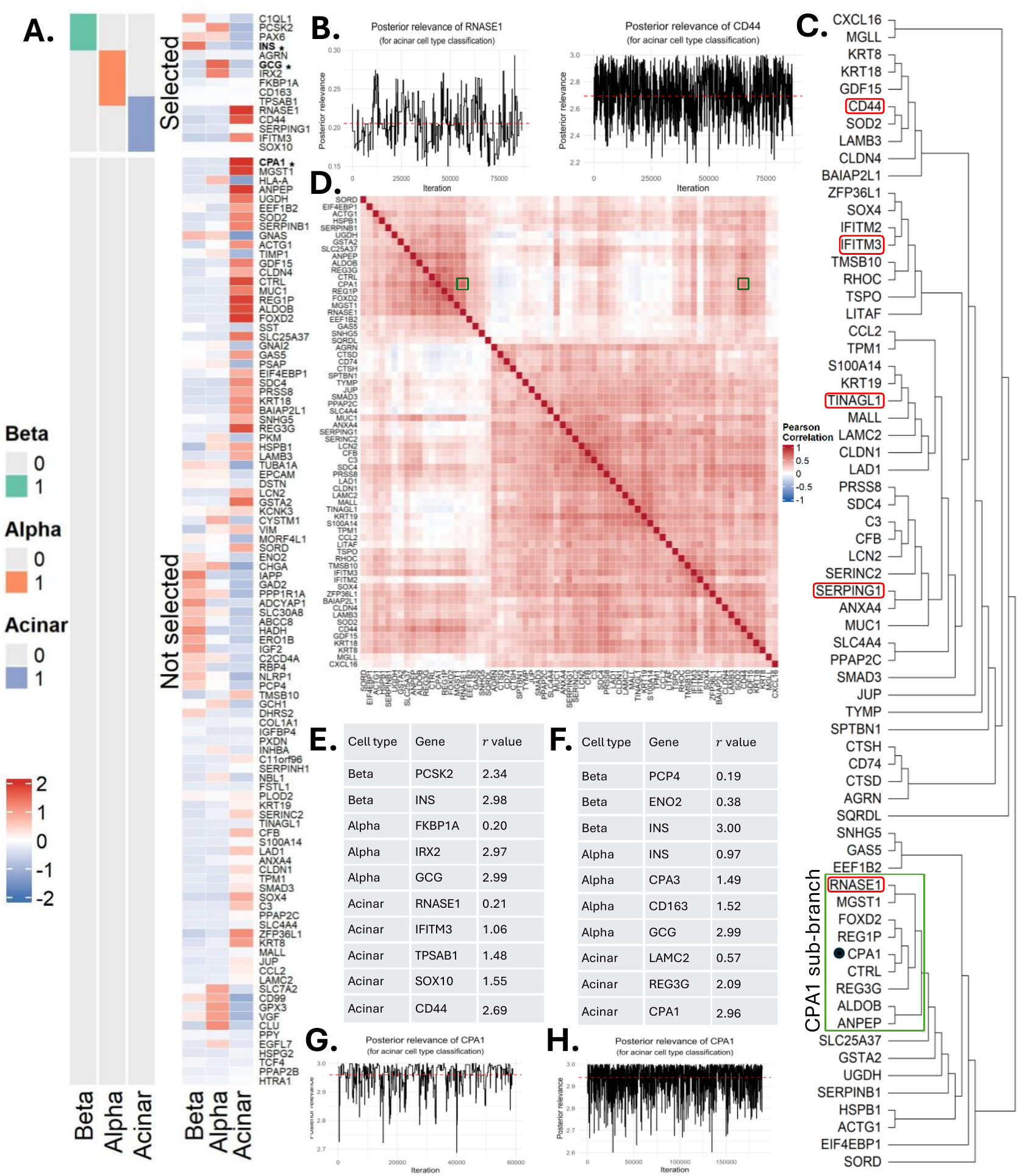
Analysis of human pancreatic cell data in Section 4.3. **A**. Heatmap of the average expression of a subset of filtered genes across three cell types. Pathologist-annotated markers for each cell type are indicated with an asterisk (*). **B**. Posterior samples of *r*_*m*_ corresponding to *CD*44 and *RNASE*1, two of the selected genes at correlation cut-off 0.75 and *FDR* < 0.05. The red dashed line represent the posterior mean of *r*_*m*_. **C**. A sub-tree of filtered genes that contain a sub-ranch of *CPA1*, denoted by a green box. Red boxes denote the marker genes from CBKMR. **D**. Correlation matrix of the genes in **C. E, F**. Posterior mean of *r*_*m*_ for the cell type markers selected at correlation-cutoff 0.75 and 0.5 respectively. **G**. Posterior samples of *r*_*m*_ corresponding to selected marker *CPA1* at correlation cut-off 0.5. **H**. Posterior samples of *r*_*m*_ corresponding to selected marker *CPA1* at correlation cut-off 0.5 and *FDR* threshold 0.99.

**Fig 7:**
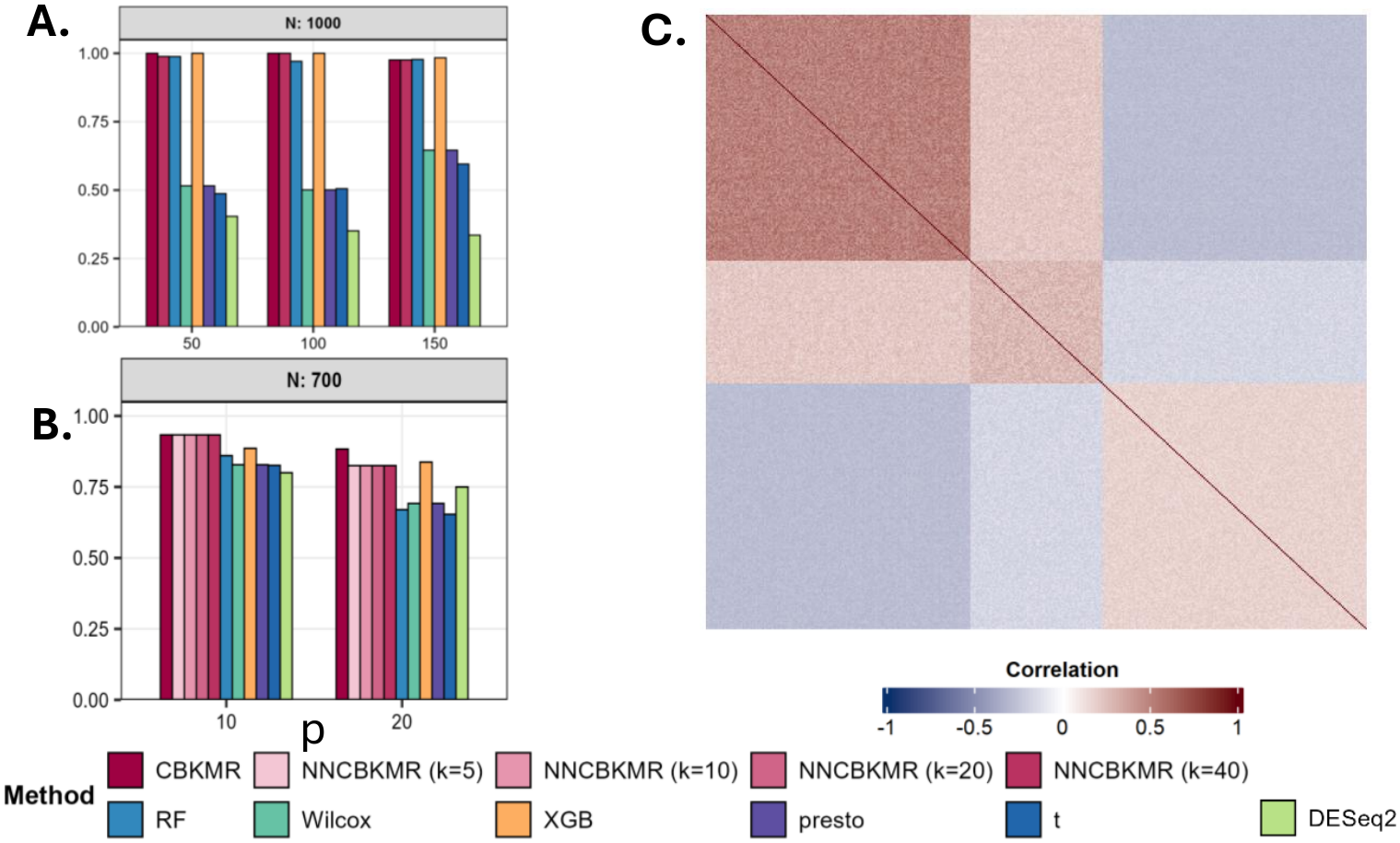
Additional simulation results for NNCBKMR in Section 3.1. A. Selection AUPRC Results corresponding to *N* = 1000 for Design 3. B. Selection AUPRC results corresponding to *N* = 700 for Design 4. C. Correlation matrix used for data generation. This correlation matrix has been used throughout the simulation studies in Section 3 for both Designs 3 and 4.

**Fig 8:**
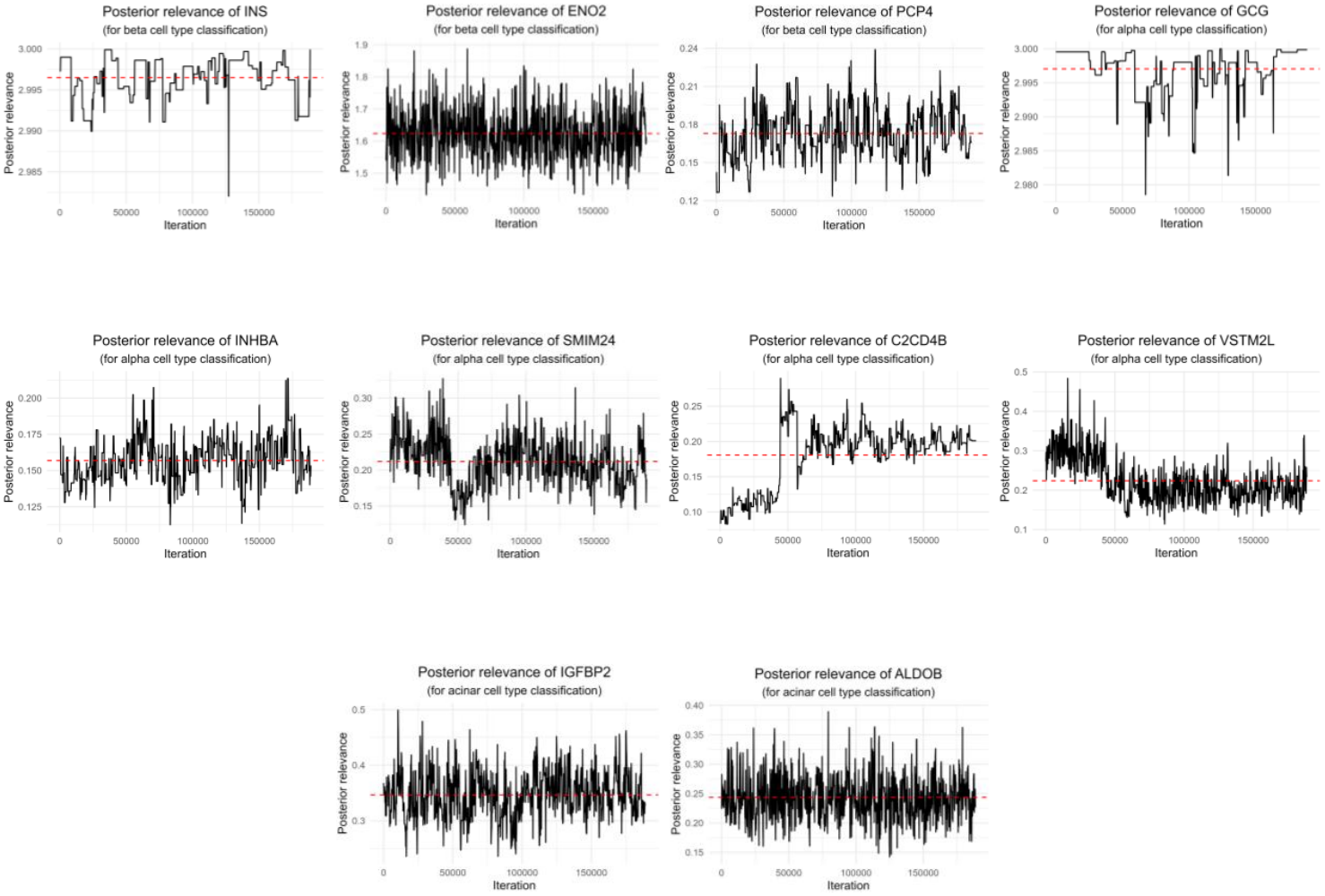
MCMC of selected genes from the pancreatic cell data in Section 4.3. *INS* and *GCG* are the pathologist annotated marekr genes for beta and alpha cell types respectively.

## 5. Conclusion

In this work, we introduce a copula-based Bayesian kernel machine regression (CBKMR), a flexible framework that separates the modeling of marginal mean structure from the kernel-induced dependence among observations. By parameterizing a Gaussian copula through an automatic relevance determination (ARD) kernel, CBKMR more reliably captures complex gene–outcome relationships and the true joint dependence structure among non-Gaussian outcomes. This improvement stems from the copula’s ability to model dependence directly at the level of the outcome distributions, rather than relying on a latent Gaussian process that induces dependence solely through the conditional mean – a construction that is well suited for continuous Gaussian responses but can misrepresent the dependence structure for discrete or otherwise non-Gaussian outcomes, as noted by Madsen (2009). To enable interpretable marker discovery, we incorporate a zero-mass spike-and-slab prior on the ARD relevance parameters, yielding intuitive variable selection through posterior inclusion probabilities (PIPs). Finally, to ensure scalability in large datasets, we develop NNCBKMR, which uses nearest-neighbor GP–based sparse Cholesky factorizations (Datta et al. (2016)) to substantially reduce computational cost.

In extensive simulation studies conducted under model misspecification (data generated from a logistic GBKMR model), logistic CBKMR consistently outperforms both logistic GBKMR and leading ensemble machine-learning methods (e.g., random forests, XG-Boost), underscoring its utility as a novel and effective marker-selection framework. Notably, NNCBKMR achieves accuracy comparable to the full CBKMR model while providing a multi-fold speedup. In applications to three scRNA-seq datasets—type 2 diabetes (T2D) islet cell types (Lawlor et al. (2017)), mouse embryo differentiation (Azzoni et al. (2021)) and regular pancreatic cells (Baron et al. (2016))—CBKMR successfully recovers nearly all pathologist-annotated marker genes in the T2D dataset and identifies biologically essential genes that separated *Ncx*1^−*/*−^ from wild-type cells in the embryo dataset. Several of these markers were down-weighted or entirely missed by feature-importance rankings from random forests and XGBoost, demonstrating the improved sensitivity of CBKMR to nonlinear and jointly informative molecular effects.

This work paves the way for broader use of copula-based kernel machine regression in settings characterized by complex, non-Gaussian dependence structures. Although CBKMR is robust to correlation among predictors, as demonstrated in our real-data analyses, a current limitation lies in the assumption of independent priors on the relevance parameters. This assumption may underrepresent the shared regulatory behavior of genes within the same biological pathway, potentially motivating future extensions that incorporate structured or group-wise dependence in the prior (Ma et al. (2024)). Additionally, further scalability may be achieved by incorporating variational inference into the NNCBKMR framework (Song and Datta (2025)), allowing fast approximate inference in large-sample settings. A final and important direction is to extend CBKMR to explicitly model spatial dependence, enabling spatially informed marker detection in emerging spatial omics datasets.

## Funding

A.C., C.M., B.N., and S.S. were supported in part by the Biostatistics Shared Resource, Hollings Cancer Center, Medical University of South Carolina (P30 CA138313). S.S. was supported by NIH R21 CA286287-01A1. A.C was supported by the American Cancer Society Institutional Research Grant: IRG-24-1290553-23-IRG. The content is solely the responsibility of the authors and does not necessarily represent the official views of the American Cancer Society, the National Cancer Institute, and the National Institutes of Health.

## APPENDIX A PROOFS OF LEMMA 2.2 AND THEOREM 2.3

### A.1. Preliminary Setup

To prove the statements in Section 2.4.1, we first fix all inverse length-scales except the *m*-th at their true values, 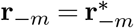 and 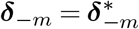. Let us define the pairwise squared-difference matrix in coordinate *m*, (*D*_*m*_)_*ij*_ = (*z*_*im*_ − *z*_*jm*_)^2^, and the partial kernel matrix

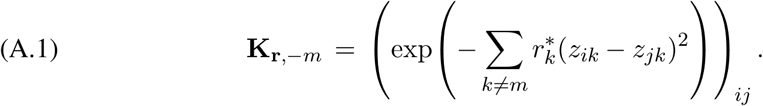

The full ARD kernel can be expressed as **K**_**r**_ = **K**_**r**,−*m*_ ⊙ **G**_*m*_(*r*_*m*_), where (*G*_*m*_(*r*_*m*_))_*ij*_ = exp(−*r*_*m*_(*z*_*im*_ − *z*_*jm*_)^2^), and ⊙ is the Hadamard (elementwise) product. The *derivative* for coordinate *m* at *r*_*m*_ = 0 can be further written as

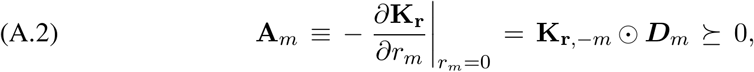

where the PSD property follows since the Hadamard product of two PSD matrices is PSD. Throughout the proofs, the following two assumptions are in force.

#### Assumption A.1

(IRLS approximation for GBKMR). For GBKMR in equation (2.1), the GP prior on ***h*** is ***h*** ∼ *N*(**0, K**_**r**_), and the IRLS working covariance at *r*_*m*_ = 0 is

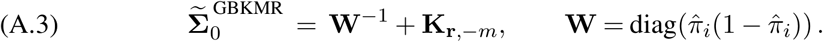

#### Assumption A.2

(CBKMR at *τ* ∈ [1*/*5, 1)). CBKMR is operated with *τ* ∈ [1*/*5, 1), estimated from the data. The copula covariance at *r*_*m*_ = 0 is

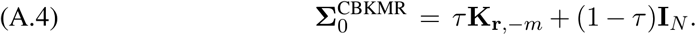

The lower bound *τ* ≥ 1*/*5 is the condition under which the Fisher information ordering in Lemma 2.2 is guaranteed; its origin is made precise in Step 1 of the proof below. In practice, CBKMR estimates *τ* close to 1, so this condition is almost always satisfied.

### A.2. Proof of Lemma 2.2

We first derive the score and Fisher information for each method, and then establish the ordering.

#### A.2.1. Score Functions and Fisher Information

##### A.2.1.1. CBKMR score

We recall the copula likelihood from equation (2.6), (A.5)

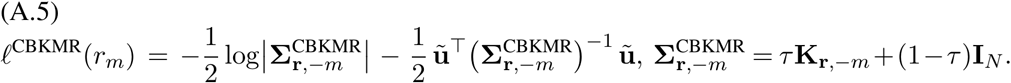

Using 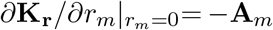 from (A.2), differentiating (A.5) at *r*_*m*_ = 0 gives (A.6)

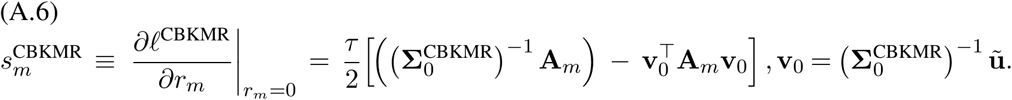

Under *H*_0_, CBKMR places 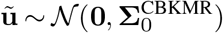. Consequently, the score statistics follow an exact Gaussian distribution with

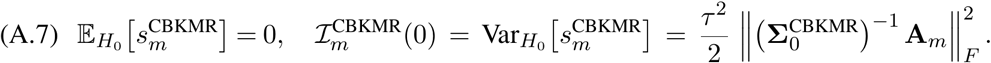

##### A.2.1.2. GBKMR score (under IRLS approximation)

Under Assumption A.1, the IRLS procedure approximates the intractable binary marginal likelihood by a Gaussian working likelihood with covariance 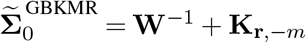. The score function at *r*_*m*_ = 0 is

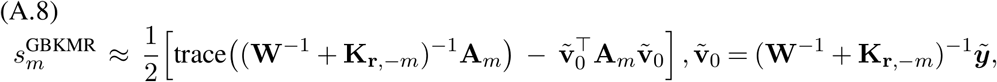

where 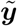 is the IRLS working response. The score function does *not* have an exact distribution: the approximation inflates the effective working covariance from **K**_**r**,−*m*_ to **W**^−1^ + **K**_**r**,−*m*_, causing score variance to exceed the curvature of the working log-likelihood. The approximate GBKMR Fisher information is

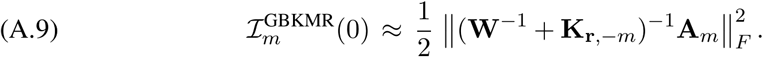

#### A.2.2. Establishing the Ordering

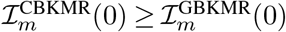, i.e., from (A.7) and (A.9),

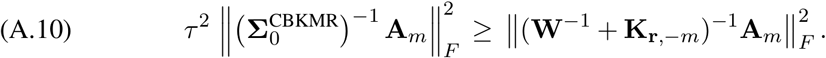

It is strarightforward to see that the above assumption is equivalent to 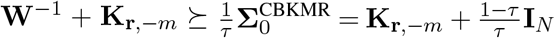. We establish this through the following steps.

**Step 1: The IRLS covariance dominates the scaled copula covariance**. The required inequality simplifies to

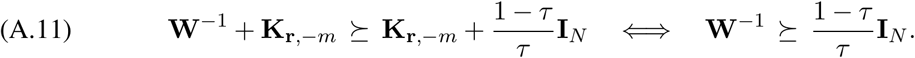

Since **W**^−1^ is diagonal with entries 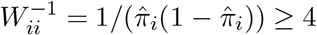,(as 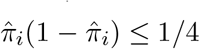for binary *y*_*i*_), condition (A.11) holds entry-wise if and only if

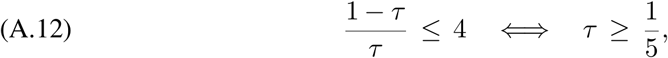

which is guaranteed by Assumption A.2. Therefore, from (A.11) and **K**_**r**,−*m*_ ⪰ 0:

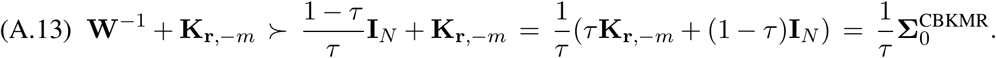

Subsequently, inverting the inequality in equation (A.13) gives us,

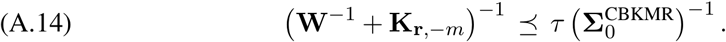

That is, the *τ* -scaled CBKMR copula precision 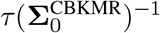 dominates the GBKMR IRLS precision (**W**^−1^ + **K**_**r**,−*m*_)^−1^ in the Loewner order.

**Step 2: The Frobenius norm ordering follows**. Since **A**_*m*_ ⪰ 0, its symmetric square root 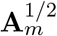 exists. Define the PSD matrices

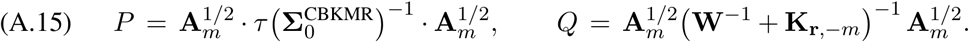

Sandwiching (A.14) by 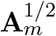 gives *Q* ⪯ *P*, i.e., *E* ≡ *P* − *Q* ⪰ 0. Writing *P* = *Q* + *E*:

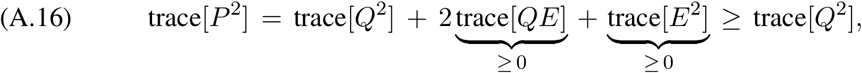

where trace 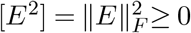 trivially, and trace[*QE*] = trace[*Q*^1*/*2^*EQ*^1*/*2^] ≥ 0 since *Q, E* ⪰ 0. Since 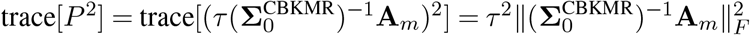 and 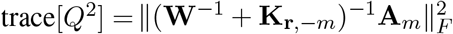, inequality (A.16) gives exactly (A.10), completing the proof of Lemma 2.2.

### A.3. Proof of Theorem 2.3

The proof proceeds in two stages: we derive the noncentrality parameters for each method, invoke Lemma 2.2 to compare them, and then establish the power ordering.

#### A.3.1. Noncentrality Parameters Under H_1_

For a local alternative with small 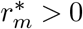, we expand 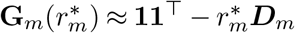, so

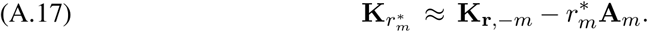

##### A.3.1.1. CBKMR noncentrality

Under *H*_1_, the working model places 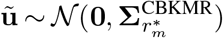 where 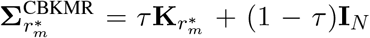. From (A.17), the shift in copula covariance is 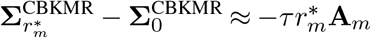. Using the score in (A.6):

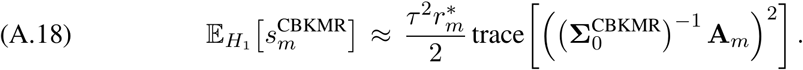

Substituting (A.18) and (A.7) into the definition of the noncentrality parameter:

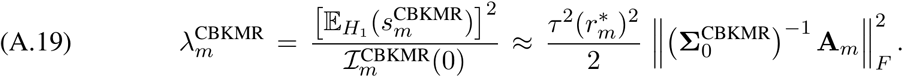

##### A.3.1.2. GBKMR noncentrality

Under Assumption A.1, the shift in the GBKMR working covariance is 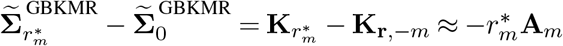. Hence, from (A.8):

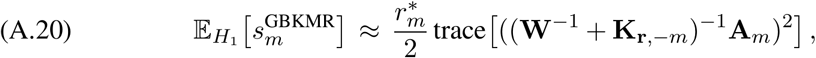

and the GBKMR noncentrality is

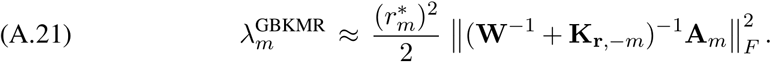

#### A.3.2. *Noncentrality Ordering*. From (A.19) and (A.21)

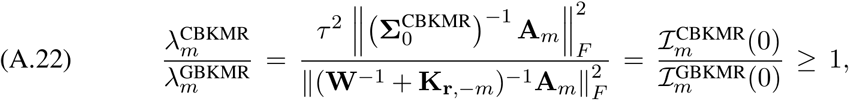

where the final inequality is precisely Lemma 2.2. Hence 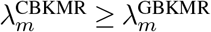

#### A.3.3. Power Ordering

The power of the level-*α* score test is

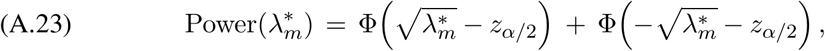

where *z*_*α/*2_= Φ^−1^(1 − *α/*2) *>* 0. We show (A.23) is strictly increasing in 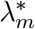. Substitute 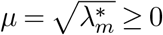 and differentiate:

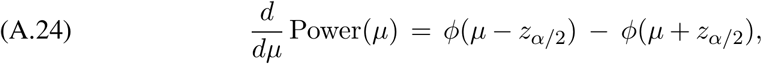

using *d*Φ(−*µ* − *z*_*α/*2_)*/dµ* = −*ϕ*(−*µ* − *z*_*α/*2_) = −*ϕ*(*µ* + *z*_*α/*2_) by symmetry of *ϕ*. Since the standard normal density 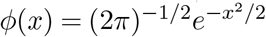 is strictly decreasing in |*x*|, and for *µ >* 0, *z*_*α/*2_*>* 0:

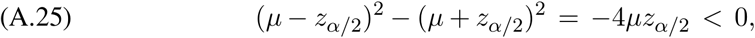

hence |*µ* − *z*_*α/*2_|*<* |*µ* + *z*_*α/*2_|, giving *ϕ*(*µ* − *z*_*α/*2_) *> ϕ*(*µ* + *z*_*α/*2_) and therefore *d* Power*/dµ >* 0. Since 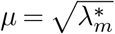 is strictly increasing in 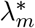, the chain rule gives 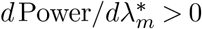 for all 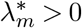. Combining this monotonicity with 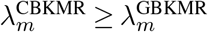 from (A.22):

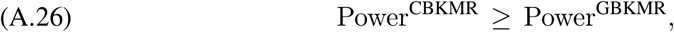

at every significance level *α* ∈ (0, 1), completing the proof of Theorem 2.3.

##### Remark A.3

(Power at boundary cases). At 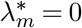 (no signal): Power(0) = 2Φ(−*z*_*α/*2_) = *α*, i.e. power equals the significance level. As 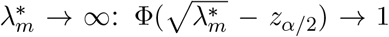 and 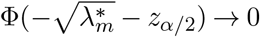, so Power → 1.

##### Remark A.4

(Robustness to the DT approximation). The distributional transform (DT) approximation introduces a per-observation variance inflation *δ*_*i*_ ≤ *π/*24 ≈ 0.131 for binary *y*_*i*_, replacing 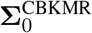 by 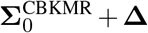 where **Δ** = diag(*δ*_*i*_) ⪰ 0. Under the DT, the Loewner condition in Step 1 becomes 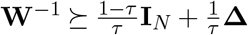, which reduces entry-wise to

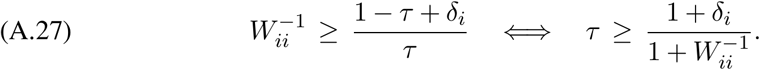

Since 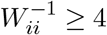 and *δ*_*i*_ ≤ *π/*24, the binding constraint is *τ* ≥ (1 + *π/*24)*/*(1 + 4) = (24 + *π*)*/*120 ≈ 0.226. Hence, the Fisher information and power orderings in Lemma 2.2 and Theorem 2.3 remain valid under the DT approximation for any *τ* ≥ (24 + *π*)*/*120 ≈ 0.226, a mild strengthening of the *τ* ≥ 1*/*5 = 0.2 condition from Assumption A.2.

## APPENDIX B PROOF OF LEMMA 2.5

We will follow Theorem 2.1 of Schäfer, Katzfuss and Owhadi (2021). for an outline of this proof. Note that, when the individual CDFs *F*_*i*_’s are continuous, there is no need for discrete transformation (DT) approximation and the density of **y** given a copula correlation matrix **Σ**_**r**_ (assuming that **y** is already permuted according to the maxmin ordering (Guinness, 2018)) can be described as,

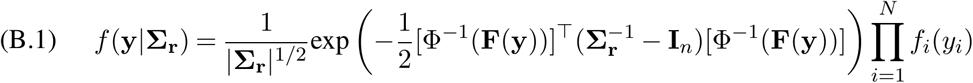

Since **Σ**_**r**_ does not contain *β*_0_ and ***β***_1_ in *f* (**y**|**Σ**_**r**_) as well as in the mathematics that follows. Once the sparsity structure *S* is obtained through kd-tree algorithm, the Kullback-Leibler (KL) divergence between the density of **y** with a copula correlation matrix **Σ**_**r**_ and the same with a copula correlation matrix 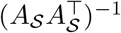 for a positive definite matrix *A*_S_ is given by,

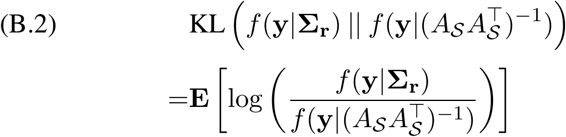

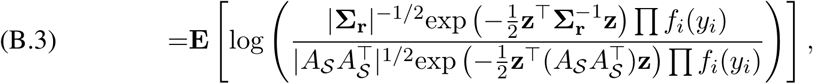

where the expectation **E** is taken with respect to the copula **y** ∼ *C*(·, **Σ**_**r**_), **z** = Φ^−1^(**F**(**y**)). The marginal densities Π *f*_*i*_(*y*_*i*_) and the identity matrices **I**_*n*_ within the quadratic forms cancel out. By applying the transformation **z** = Φ^−1^(**F**(**y**)), the expression simplifies to the KL divergence between two centered multivariate Gaussian distributions:

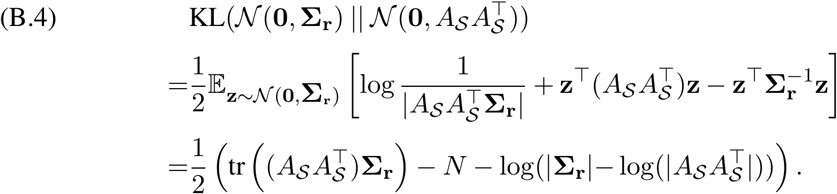

Since *A*_S_ is sparse lower triangular matrix, we can directly argue following Theorem 2.1 of Schäfer, Katzfuss and Owhadi (2021) that the KL term will be minimized when *A*_S_ = **U**_S_, **U**_S_ is described as in Lemma 2.5.

## APPENDIX C

ADDITIONAL SUMMARIES FROM SIMULATION EXPERIMENTS

In this section, we provide additional summaries from the simulation experiments in Section 3.

## SUPPLEMENTARY MATERIAL

### Additional data analysis results and traceplots

Contains discussions on correlation filtering, implementing CBKMR on scDesign3.

### Reproducible codes

Contains CBKMR package and reproducible codes for the analyses

## Notes

### Competing Interest Statement

The authors have declared no competing interest.

### Summary of Updates

Addition of new theorems and simulation results

