## Supplementary Material for "CBKMR: A Copula-Based Bayesian Kernel Machine Regression Framework for Optimal Marker Detection in Omics Data"

**1. Correlation filtering .** In Section 4.2 and 4.3 of the manuscript, we have shown that a moderately large strong correlation among the covariates may produce suboptimal results during marker identification task. This is because of the fact that allowing a strong correlation introduces practically non-identifiability among the covariates. To understand this, let us consider a simple linear regression model with a continuous response $y$ as $y = a_0 + a_1x_1 + a_2x_2 + a_3x_3 + \epsilon$, where $x_{i,1\le i\le 3}$ are the covariates. Now, if $x_1 = c + dx_2,\ d \neq 0$, i.e., cor$(x_1, x_2) = 1$, then $y = a_0 + a_1x_1 + a_2x_2 + a_3x_3 + \epsilon = (a_0 + a_1c) + 0x_1 + (a_2 + a_1d)x_2 + a_3x_3 + \epsilon$. Therefore, no single value of the tuple $(a_0, a_1, a_2)$ can satisfy the equation. This phenomenon called multicollinearity (Chan et al., 2022). In practice, when the measurement error is moderate, the estimated values of the coefficients in linear regression can under or overestimate the true values. To illustrate this, we simulated a synthetic dataset comprising of 100 observations $y_1, \ldots, y_{100}$ with 30 feature vectors $z_{i1}, \ldots, z_{i30}$ for each observation. Additionally, we introduced a permissive correlation of 0.75 among $z_{i2}, \ldots, z_{i12}$. Noticably, this correlation cutoff was used in the data analysis in Section 4. Finally, we considered three covariates as significant in the data generating model, with two covariates from the correlated block: $y_i = 1 + 0.5z_{i1} + z_{i2} + 2z_{i12} + \epsilon_i$, where $\epsilon_i \sim \mathcal{N}(0, 1)$. A template of the R code is given below.

```
N <- 100; p = 30

Sigma_Z <- diag(1, N) #feature covariance, assuming the features to be independent across samples

Z <- list()
dframe <- NULL

seed <- 135
set.seed(seed)
for(i in 1:p){
  Z[[i]] <-  c(mvnfast::rmvn(n = 1, mu = rep(0, N), sigma = Sigma_Z))
}
Z <- do.call(cbind, Z)                              # final feature matrix, N x p
Sigma_corr <- matrix(0.75, 10, 10)

diag(Sigma_corr) <- 1
datamat <- mvnfast::rmvn(n = N, mu = rep(0, 10), sigma = Sigma_corr)
```

```

Z[ , 2:11] = datamat

###################################################################
########## Linear model -- frequentist optimization ##############
###################################################################
ytrue = as.numeric(1 + 0.5 * Z[ , 1] + 1 * Z[ , 2] + 2 * Z[ , 12])

set.seed(seed)
y = ytrue + rnorm(N, sd = 1.0)
# 1. Fit the standard linear model
fit_freq <- lm(y ~ Z)

# 2. Look at the summary
summary_test <- summary(fit_freq)
summary_test

# Call:
#   lm(formula = y ~ Z)
#
# Residuals:
#    Min       1Q   Median       3Q      Max
# -1.60667 -0.41249  0.02536  0.42214  1.67000
#
# Coefficients:
#              Estimate  Std. Error  t value  Pr(>|t|)
# (Intercept)  0.938268   0.097803   9.593 2.51e-14 ***
# Z1           0.460642   0.096061   4.795 9.03e-06 ***
# Z2           1.339468   0.192643   6.953 1.62e-09 ***
# Z3           0.201134   0.167102   1.204   0.2328
# Z4          -0.067315   0.187525  -0.359   0.7207
# Z5          -0.429401   0.174459  -2.461   0.0163 *
# Z6          -0.094666   0.163498  -0.579   0.5645
# Z7           0.014294   0.165634   0.086   0.9315
# Z8          -0.008207   0.163560  -0.050   0.9601
# Z9          -0.004824   0.150689  -0.032   0.9746
# Z10         -0.453495   0.206890  -2.192   0.0318 *
# Z11          0.207648   0.177329   1.171   0.2456
# Z12          2.229843   0.111844  19.937  < 2e-16 ***
# Z13          0.090627   0.088873   1.020   0.3114
# Z14         -0.081974   0.105621  -0.776   0.4403
# Z15         -0.007037   0.086905  -0.081   0.9357
# Z16         -0.211412   0.091355  -2.314   0.0236 *
# Z17          0.109324   0.097522   1.121   0.2662
# Z18         -0.002550   0.087120  -0.029   0.9767
# Z19          0.069882   0.106510   0.656   0.5139
# Z20          0.114230   0.091661   1.246   0.2169
# Z21          0.035704   0.096189   0.371   0.7116
# Z22          0.015728   0.101962   0.154   0.8779
# Z23          0.012016   0.112102   0.107   0.9150
# Z24         -0.125193   0.104243  -1.201   0.2339
# Z25          0.074804   0.092141   0.812   0.4197
# Z26         -0.075234   0.110605  -0.680   0.4987
# Z27          0.077251   0.083520   0.925   0.3582
```

```
# Z28          -0.080517   0.093170  -0.864   0.3905
# Z29          -0.085316   0.085162  -1.002   0.3199
# Z30          -0.020590   0.089152  -0.231   0.8180
# ---
#   Signif. codes:  0 '***' 0.001 '**' 0.01 '*' 0.05 '.' 0.1 ' ' 1
#
# Residual standard error: 0.7968 on 69 degrees of freedom
# Multiple R-squared:  0.9072,   Adjusted R-squared:  0.8669
# F-statistic: 22.49 on 30 and 69 DF,  p-value: < 2.2e-16
```

The linear model identifies the three true predictors, $z_{\cdot 1}$, $z_{\cdot 2}$, and $z_{\cdot 12}$, as statistically significant. However, the estimated coefficients of $z_{\cdot 2}$ and $z_{\cdot 12}$, 1.34 and 2.23, respectively, deviate from their true values of 1 and 2. Moreover, several inactive predictors receive nonnegligible coefficient estimates, with $z_{\cdot 5}$ and $z_{\cdot 10}$ incorrectly identified as significant. Because these variables belong to the correlated block containing $z_{\cdot 2}$, the signal associated with $z_{\cdot 2}$ may be distributed across multiple correlated predictors, producing unstable coefficient estimates and spurious selections. This behavior is consistent with the observed variance inflation factors, which range from approximately 3.16 to 6.23 within the correlated block.

A similar identifiability challenge can arise in kernel-based models such as CBKMR. The relevance parameters $r_m$ determine the contribution of each feature to the kernel function, and when features are strongly correlated, distinct configurations of $\{r_m\}$ may produce nearly identical kernels, creating a geometric identifiability issue that affects variable selection. Consequently, removing highly correlated features before model fitting is a well-established strategy for reducing the effective feature space of multivariate nonlinear models (Wang et al., 2005; Perez-Riverol et al., 2017; Abdulrauf Sharifai and Zainol, 2020).

However, CBKMR is notably more tolerant of correlation among predictors than existing per-gene testing methods such as Wilcoxon, Presto, or DESeq2. In Section 3 of the main manuscript, we demonstrate through extensive simulation that CBKMR achieves superior variable selection performance under a moderate predictor correlation of 0.7. Critically, unlike linear regression where correlated inactive predictors may receive significant coefficient estimates and produce false positives, the simulation results show that CBKMR produces on average fewer false-positive selections even under moderate correlation. The false-positive behavior observed in the linear model therefore does not carry over to CBKMR in our simulation settings. In practice, only genes exhibiting near-perfect redundancy, specifically those with pairwise correlation exceeding 0.75, require removal prior to fitting CBKMR.

**2. Discussion on `scDesign3` simulated data.** In recent years, several software packages have been developed to generate realistic single-cell transcriptomic datasets (Zappia, Phipson and Oshlack, 2017; Dibaeinia and Sinha, 2020; Song et al., 2024; Luo et al., 2024). Among these, scDesign3 (Song et al., 2024) provides a comprehensive framework for simulating scRNA-seq and spatial transcriptomic data conditioned on cell-type labels, developmental lineages, and batch indicators. Critically, however, these frameworks follow a top-down generative structure: cell-type labels are fixed a priori, and gene expression profiles are subsequently sampled conditional on those labels. Under this generative structure, the simulator encodes which genes are differentially expressed across cell types, but does not specify which genes are predictive of cell-type identity in the supervised sense. Consequently, such pipelines do not provide a well-defined ground truth for evaluating gene-level predictive importance, making it difficult to benchmark methods such as CBKMR that aim to identify genes whose expression is informative of cellular phenotype. To further illustrate this, we considered the sce_filteredExpr10_Zhengmix4eq dataset used in `scDesign3`. Following the software's copula-based framework, we simulated a dataset with $N = 800$

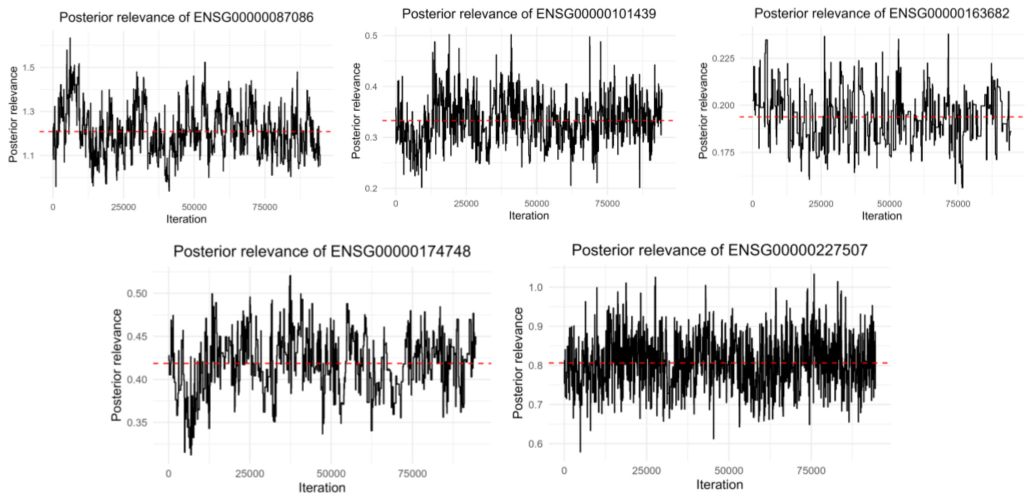


Fig 1: CBKMR results from the simulated dataset using scDesign3.

cells and $p = 400$ genes, restricting the analysis to B cells and CD14 monocytes. Correlation filtering was subsequently applied as described in Section 1, removing genes with pairwise correlation exceeding 0.9 with any retained gene. Figure 1 presents MCMC diagnostics for five selected genes with average posterior $r_m > 0.1$. Among these, ENSG00000087086, ENSG00000101439, and ENSG00000227507 were differentially expressed between the two cell types. Since these simulation frameworks do not define a ground truth for cell-type-predictive gene identification, the remaining genes cannot be evaluated in terms of true positive recovery.

Despite this limitation in ground truth specification, realistic simulators such as scDesign3 serve a complementary purpose: they faithfully reproduce empirical properties of scRNA-seq data, including sparsity and gene-gene correlation structure, which are difficult to capture through purely parametric designs. The sparsity of the above simulated dataset was estimated at approximately 34%, consistent with our parametric simulation settings. Informed by these empirical characteristics, we designed our own simulation framework that preserves similar levels of sparsity and permits moderate pairwise gene correlations of up to 0.6. Critically, our design explicitly assigns ground-truth cell-type-predictive genes, with binary cell-type labels generated from gene expression features via a Bernoulli distribution, enabling rigorous evaluation of marker recovery. The complete data-generating procedure is implemented in the function `generate_sparse_nb_data`, made publicly available on GitHub.
